# Interplay of Long- and Short-term Synaptic Plasticity in a Spiking Network Model of Rat’s Episodic Memory

**DOI:** 10.1101/2024.06.21.598805

**Authors:** N. Chrysanthidis, F. Fiebig, A. Lansner, P. Herman

## Abstract

We investigated the interaction of long-term episodic processes with effects of short-term dynamics of recency. This work takes inspiration from a seminal experimental work involving an odor-in-context association task conducted on rats (Panoz-Brown et al., 2016). In the experimental task, rats were presented with odor pairs in two arenas serving as old or new contexts for specific odors-items. Rats were rewarded for selecting the odor that was new to the current context. New odor items were deliberately presented with higher recency relative to old items, so that episodic memory was put in conflict with non-episodic recency effects. To study our hypothesis about the major role of synaptic interplay of long- and short-term plasticity phenomena in explaining rats’ performance in such episodic memory tasks, we built a computational spiking model consisting of two reciprocally connected networks that stored contextual and odor information as consolidated and distributed memory patterns (cell assemblies). We induced context-item coupling between the two networks using Bayesian-Hebbian plasticity with eligibility traces to account for reward based learning. We first reproduced quantitatively and explained mechanistically the findings of the experimental study, and further simulated alternative tasks, e.g. where old odor items were instead encoded with higher recency, thus synergistically confounding episodic memory with effects of recency. Our model predicted that higher recency of old items enhances item-in-context memory by boosting the activations of old items resulting in further enhancement of memory performance. We argue that the model offers a computational framework for studying behavioral implications of the synaptic underpinning of different memory effects in experimental episodic memory paradigms.

**Significance Statement:** An important aspect of computational modeling is its ability to bridge spatial scales. Our cortical memory model represents a novel computational attempt to unravel neural and synaptic processes with mesoscopic manifestations underpinning the complex effects of short-term memory dynamics on episodic memory recall. We consider the quantitative match with Panoz-Brown et al.’s (2016) experimental findings, obtained in a detailed spiking network model constrained by available biological data, a significant step towards bridging the gap between behavioral correlates of episodic memory and synaptic mechanisms. Our findings and additional predictions on a suite of different episodic memory tasks invite further experimental examination.

## Introduction

Episodic memory refers to an ability to recall past experiences. The uniqueness of these memories lies in their specific environmental context, as they are memorized in particular spatial locations at a given time (Yonelinas et al., 2019). Despite the multitude of past experiences, often sharing some contextual similarity, they can be vividly distinguished due to the specificity of the overall context with its episodic, typically both spatial and temporal, characteristics. Consequently, we can usually reliably order such long-term episodic memories in time (Tulving 1972, 1985). It is less clear however how non-episodic short-term memory phenomena, inevitably accompanying episodic recall scenarios for more recently encoded memories, affect the episodic memory capability. After all, the contextual binding that underlies episodic memory should be unique to specific events, experiences with their temporal footprint. To date, the effect of recency, sometimes confounded with familiarity (Zhang et al., 2023), has been only sporadically examined in experimental studies concerned with episodic recall. In consequence, partly due to a reductionist approach to computational modeling of episodic memory phenomena, there is no emerging hypothesis about the neural and synaptic mechanisms maintaining the dynamic interaction between long- and short-term memory processes. Panoz-Brown et al.’s (2016) seminal behavioral study on episodic memory in rats revealed some new vital insights largely owing to their novel experimental design. Namely, they adapted an odor-span task involving a sequence of recently experienced, yet overall familiar, odors to an episodic memory test with distinct environmental contexts – arenas where the odors were presented. Rats were rewarded for selectively responding to only those odors that were new to any given arena (new-in-context stimuli). To directly contrast recency and context-dependent (episodic) memory effects, new-in-context odors were typically presented more recently than odors previously encountered in the given context (old-in-context) prior to pairwise (“new” vs. “old”) odor Memory Assessment. The task was arranged so that rats to be successful would have to overcome the short-term memory recency bias of new items and rely on an episodic association encoded earlier between a given old-in-context odor and the contextual arena. In other words, there was a competition between the recency of short-term odor memory and long-term episodic item-in-context (odor in an arena) memory binding. Rats turned out to overcome this recency bias and reliably performed episodic recall to successfully complete the task and claim reward, even for retention intervals reaching 45 minutes. Inspired by the Panoz-Brown et al.’s (2016) study, we built a computational spiking neural network model to investigate neural mechanisms underlying the interplay between episodic memory and short-term memory effects of recency at a mesoscopic network level. In other words, our ambition was to provide novel mechanistic insights into these complex and scarcely examined synergistic memory phenomena by, first, explaining the behavioral results reported by Panoz-Brown et al. (2016) as the emergent network effect of local synaptic plasticity phenomena at varying time scales and, second, generating testable predictions for behavioral outcomes in modified experimental paradigms.

Our biologically detailed spiking neural network model consists of two modular attractor memory networks that store contextual information (2 contexts) and odor items (16 odors), respectively, as distributed long-term (consolidated and thus familiar) memory patterns. Familiarity reflects recognition of the embedded items without any retrieval of its associated contextual information (Merkow et al., 2015). We simulated the process of encoding episodic memories in line with the experimental task design proposed by Panoz-Brown et al. (2016) as associative between-network connections binding odor and context memory items shaped by a range of synaptic plasticity effects including Hebbian plasticity, synaptic depression and augmentation as well as intrinsic plasticity (neural excitability) and spike frequency adaptation. Importantly, we accounted for the reward effect since it was relevant in the experiments not only as an incentive for rats to perform the task but predominantly as a rapid learning cue in this complex continual learning paradigm. To that end, we employed eligibility traces in the framework of our Bayesian-Hebbian synaptic learning rule and upregulated associative plasticity upon successful odor-in-context recall. We simulated recall as a discriminative process between neural activities attributed to competing (old vs. new) odor memory patterns presented as a pair of odor network stimuli in rapid succession with the simultaneous contextual cue active in the background. We demonstrated how the combination of different synaptic processes contributed to the observed item-in-context memory. We also simulated an alternative version of the original task, where the order of odor presentation was switched so that old items-in-context were more recently encoded than new items, to quantify the memory recall enhancement due to the synergistic contribution of episodic and short-term memory effects. Finally, we tested the resistance of episodic memory to interference by simulating yet other challenging task variations, introducing additional contextual information (extra context) or altering the task structure by violating the balanced odor item training scheme (repeating some items more times than others).

## Results

Considerable experimental effort has been invested in demonstrating episodic memory in rats using item-in-context paradigms. In such paradigms, rats are trained to recognize items across multiple contexts (Panoz-Brown et al., 2016; O’Brien & Sutherland, 2007; Lesburguères et al., 2017; Bevins and Besheer, 2006). A crucial challenge lies in effectively dissociating episodic encoding from short-term effects, such as recency. Panoz-Brown et al. (2016) devised an item-in-context task that allows these processes to interact and compete, and thus provide valuable evidence for their implications on episodic memory performance in rats. Here we aimed to explain behavioral implications of different memory phenomena mechanistically in terms of their underlying neural and synaptic basis. We hypothesized that the interplay of different synaptic plasticity mechanisms at varying time scales is reflected in the functional connectivity of the learned network (synaptic weights) and manifests itself in the network activity (firing rates, see Methods), which in turn should help us interpret the memory performance reported in the behavioral experiment. More broadly, we wanted to address a general question of how episodic recall is subject to short-term memory phenomena, ubiquitous in real-world scenarios, at the level of network dynamics driven by synaptic plasticity mechanisms. To this end we employed a computational model consisting of two inter-connected spiking neural networks storing odor-item and context-arena memories, respectively. Accordingly, before simulations of the experimental trial blocks, long-term (well established, consolidated) item and context memory patterns were first embedded by means of prior Bayesian-Hebbian learning with multiple epochs and a long plasticity time constant. The resulting within-network attractor projections (within-network connectivity, solid red lines, Fig. 1A) remained then fixed throughout the simulated task. Contrarily, bidirectional associative connections between item-context pairs (between-network connectivity, dashed red lines in Fig. 1A) were plastic during the simulations of the experimental block, i.e. subject to on-line Bayesian-Hebbian learning with long episodic plasticity time constant and to other known short-term plasticity mechanisms (Erickson et al., 2010; Lisman, 2017).

**Figure 1:**
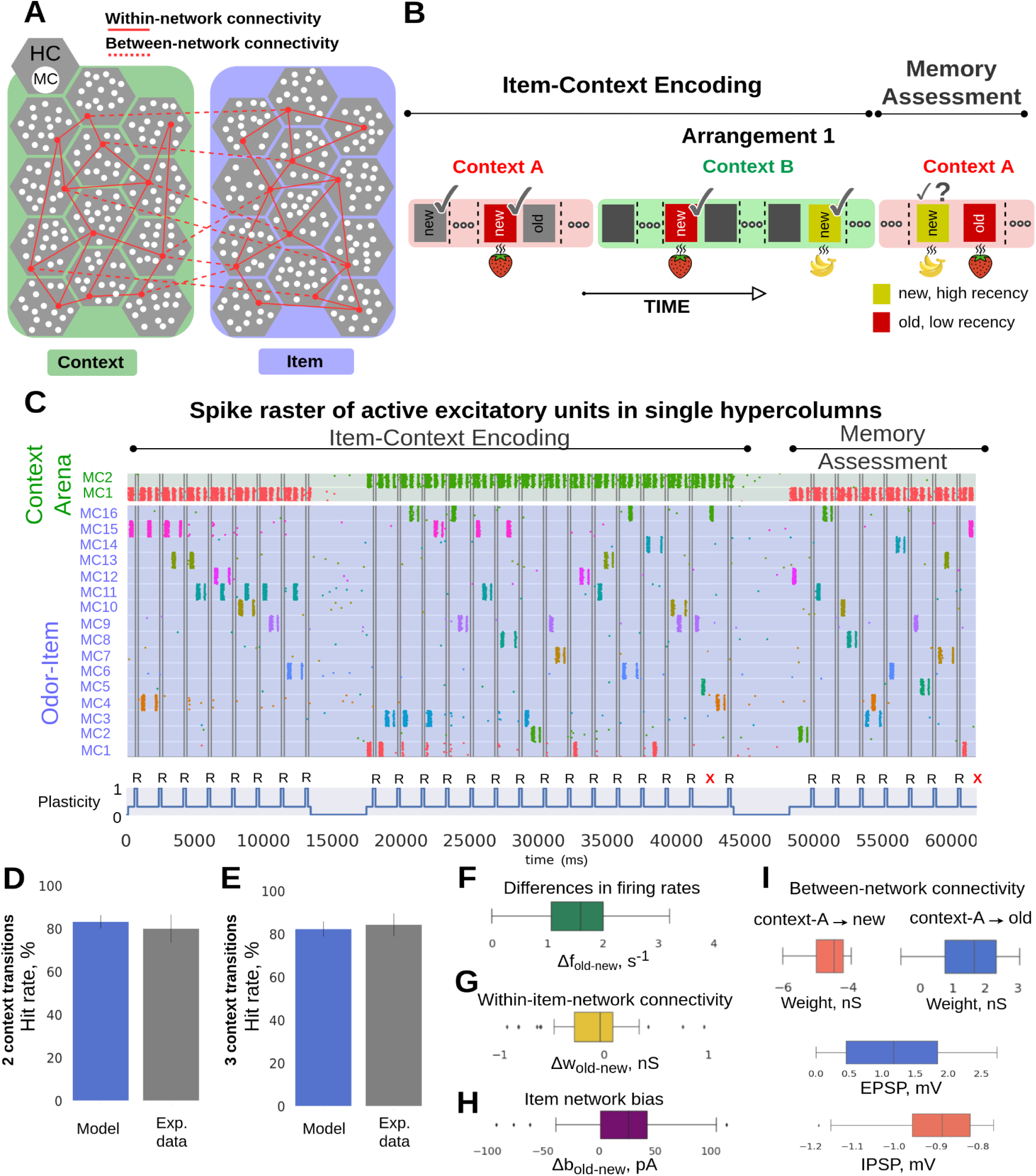
Item-in-context memory network model relying on associative episodic binding. **A**, Graphic illustration of the Item (green) and Context (blue) networks. Attractor connections (solid red) represent within-network connectivity across hypercolumns (HCs) in the same network, while associative binding refers to the plastic connections between Item and Context networks (dashed red). **B**, Task structure: odors were presented across two contexts in the simulated episodic memory task. Schematic of the two-context-transition task displaying pairs of new-old odors (depicted as rectangles with unique colors) in a given context [cf. Fig. 1B in Panoz-Brown et al., (2016)]. Only the new items-in-context were rewarded (✓ symbol in the schematic denotes reward) when selected (a 50 ms stimulation of the selected odor preceded the reward phase, representing a final odor sniff before the reward). Once a new item was presented it was considered as old for the subsequent trials in the given context (as a trial we defined a stimulation of a pair of new- and old-in-context items). Items were stimulated for the first time in context A, half of the total 16 items were presented and rewarded in context A. After the context transition all the 16 items were presented in random pairs in context B. Finally, Memory Assessment was made in context A, where we presented the remaining half of the items that had not been presented in context A (new items), and paired them randomly with old items (pairs of odors were different throughout the task). Context representations were constantly activated while cueing pairs of new-old items for 250 ms each. In the Memory Assessment block, pairs of new-old items followed the Arrangement 1 (new items were encoded more recently than old ones). Here, we show only 4 out of 16 items stimulated during the task (blue and yellow items illustrate Arrangement 1). The presence of the additional items, which are not shown is indicated as “…”. **C**, Spike raster of pyramidal neurons in HC1 of the Item and Context networks simulating the episodic memory task described in (B). Item and context memory patterns are represented by the activation of a unique set of minicolumns (MCs) in their network. Each item or context was assigned with a unique color. While context representations were persistently cued we activated new and old items-in-context during trials. Plasticity of the associative binding between Item and Context networks was modulated during item presentation and rewarded accordingly (bottom subplot, R symbol in the schematic denotes reward, and **X** symbol, in red, indicates a failed trial). **D**-**E**, The model discriminates between new- and old-in-context items with performance quantitatively matching Panoz-Brown et al.’s (2016) behavioral results in Experiment 1 (D) and 2 (E), respectively. Error bars represent SDs derived from the Bernoulli distributions for the probabilities of success (hit) across all trials (scaled to %), and for original experimental results - data is shown as mean +1 SEM across rats. **F**, Boxplot of the differences in average firing rates between pairs of old vs. new items, Δf_old-new_. **G**, Distribution of the differences in average within-network connectivity between pairs of old vs. new items, Δw_old-new_, (within-network connectivity includes short-term plasticity mechanisms combined with long-term Hebbian component: AMPA and slower NMDA receptor mediated weights). **H**, Distribution of the differences in average intrinsic excitability between pairs of old vs. new items, Δbias_old-new_. **I**, Weight distribution of the episodic associative binding prior to the Memory Assessment part of the task. The distributions display the means of the learned synaptic weights (AMPA and slower NMDA receptor mediated weights, see Table 1) between context A and new (blue), or old (red) items. The inset displays the excitatory and inhibitory postsynaptic potentials (at a biological plausible range, Wang et al., 2006) of the corresponding weights distributions, which also account for the multiplicative effect of synaptic augmentation.

**Table 1:** Neuron model and synaptic parameters.

| Neuron model parameter | Symbol | Value | BCPNN parameter | Symbol | Value |
| --- | --- | --- | --- | --- | --- |
| Adaptation current | $b$ | 86 pA | BCPNN AMPA gain | $w_{gain}^{AMPA}$ | 0.33 nS |
| Adaptation decay time constant | $\tau_{lw}$ | 280 ms | BCPNN NMDA gain | $w_{gain}^{NMDA}$ | 0.03 nS |
| Membrane capacitance | $C_m$ | 280 pF | BCPNN bias current gain | $\beta_{gain}$ | 40 pA |
| Leak reversal potential | $E_L$ | -70.6 mV | BCPNN lowest spiking rate | $f_{min}$ | 0.2 Hz |
| Leak conductance | $g_L$ | 14 nS | BCPNN highest spiking rate | $f_{max}$ | 25 Hz |
| Upstroke slope factor | $\Delta_T$ | 3 mV | BCPNN lowest probability | $\epsilon$ | 0.0026 |
| Spike threshold | $V_t$ | -55 mV | P trace time constant | $\tau_p$ | 30 s |
| Spike reset potential | $V_r$ | -60 mV | E trace time constant | $\tau_e$ | 500 ms |
| Refractory period | $\tau_{ref}$ | 5 ms | AMPA Z trace time constant | $\tau_Z^{AMPA}$ | 5 ms |
| | | | NMDA Z trace time constant | $\tau_Z^{NMDA}$ | 100 ms |
| | | | Regular plasticity | $\kappa_{normal}$ | 0.3 |
| | | | Modulated plasticity | $\kappa_{reward}$ | 1 |
| Receptor parameter | Symbol | Value | Short-term plasticity parameter | Symbol | Value |
| AMPA synaptic time constant | $\tau^{AMPA}$ | 5 ms | Utilization factor | $U$ | 0.2 |
| NMDA synaptic time constant | $\tau^{NMDA}$ | 100 ms | Augmentation decay time constant | $\tau_A$ | 5 s |
| GABA synaptic time constant | $\tau^{GABA}$ | 5 ms | Depression decay time constant | $\tau_D$ | 280 ms |
| AMPA reversal potential | $E^{AMPA}$ | 0 mV | | | |
| NMDA reversal potential | $E^{NMDA}$ | 0 mV | | | |
| GABA reversal potential | $E^{GABA}$ | -75 mV | | | |

In fact, we used the same dual network model that was initially built to propose and assess a Bayesian-Hebbian hypothesis about synaptic and network mechanisms underlying semantization of episodic memory, i.e. transformation of episodic memories to more abstract semantic representations (Chrysanthidis et al., 2022). The model reflects a wide range of biological constraints and operates on behavioral time scales under constrained network connectivity with plausible postsynaptic potentials, spiking activities, and other biophysical parameters (see Methods).

### Episodic memory contra recency effects: The control task design (Arrangement 1)

We first used the model to simulate Panoz-Brown et al.’s (2016) base experimental setup, where rats were exposed to a rapid presentation of several odors across two arenas (A,B) serving as contexts for odor items (A ➞ B ➞ A, Experiment 1, Fig. 1B; Symbol “➞” indicates a context transition). Our simulations followed the item-context association protocol adopted from the two-context-transition task denoted as “Experiment 1” in Panoz-Brown et al.’s (2016) study. It consisted of two main experimental blocks: Item-Context Encoding and Memory Assessment. In the first block, 8 odor items were presented in context A, followed by all 16 odor items presented in context B (one context transition, A ➞ B). So half of the items presented in context B were previously encoded in context A. In the experiment, odors were always presented in pairs, new- vs. old-in-context items, and rats responded by selecting one of them as new, which marked an individual trial. In the model, we cued corresponding memory item patterns in the Item network in short succession (inter-stimulus period of 250 ms, see Methods) to simulate the serial process of first recognising one odor then the other in each pair (as a result of sniffing). Simultaneously, we cued the respective memory pattern in the Context network to account for contextual information, which resulted in cross-network binding of the cell assemblies representing odor items and contexts via associative Hebbian plasticity.

In the second block, referred to as Memory Assessment, every remaining odor (8 out of 16), new to context A, was presented in a pair with another randomly selected odor that was considered at the presentation time as already encoded in that context, i.e. an old-in-context item. To reiterate, in this original task design proposed by Panoz-Brown et al. (2016) old-in-context items featured lower recency (Fig. 1B, Arrangement 1, old items were always presented earlier than new items prior to Memory Assessment block), so that correct retrieval of items had to entirely depend on the contextual association. Recency might increase the sense of familiarity of an item, thereby potentially confusing rats. Therefore, in that arrangement, context-dependent episodic memory recall was put into conflict with the effect of memory recency (Fig. 1B, Arrangement 1). Old- and new-in-context items were cued only once during the Memory Assessment block. This is in contrast with the aforementioned Item-Context Encoding block where different items could be repeated multiple times and thus old-in-context items had always higher recency compared to new-in-context items (here: old/new relative the given context in the Item-Context Encoding block) as the old items were encoded in the same context before the new item was presented. Odor pairs were different between Item-Context Encoding block and Memory Assessment, and also randomized across simulations. Accordingly, behavioral data in the experimental study and simulated data of the model performance here were examined only during the Memory Assessment block.

In Figure 1C we illustrate an exemplary spike raster of active pyramidal neurons in one of the network hypercolumns (see Methods) of both the Item and the Context network obtained in a simulation of the entire experimental session. The bottom of Figure 1C depicts the associative plasticity gain (item-context binding) modulation. It accounts for a reward signal (Fig. 1C, “R”: reward) that in line with the original experiment follows each successful odor choice (a continual learning scenario). The reward implementation uses synaptic eligibility traces and temporarily boosts associative plasticity gain from the baseline level, κ_normal_ (Table 1), during item presentation to the elevated κ_reward_ (Table 1). The odor choice (old- vs new-in-context) itself was made based on a comparison between average firing rates elicited by the excitatory units corresponding to the two stimulated (competing) item patterns in each pair (see Methods).

By tuning stimulus-related parameters (i.e., strength of simulations and background noise excitation) of our earlier model on item-context episodic memory binding (Chrysanthidis et al., 2022), we obtained task performance comparable to the original experimental data (Fig. 1D,E; model data: mean=83.21, SD=3.12, n=143, mean represents the total number of successes across all n-trials [each trial tests one old-new pair], SD derived from the Bernoulli distributions for the probabilities of successes across all n-trials, n corresponds to simulated old-new pairs during Memory Assessment, and experimental data: mean≈80, SD≈6.5, mean reflects the averaged performance of rats in 9 sessions, combining the initial and terminal sessions, and SD reflects the averaged standard error of the mean (SEM) across rats for the combined initial and terminal sessions) for the two-context-transition task (Experiment 1, see Panoz-Brown et al., 2016). The high odor item recall performance of the model originates from considerably stronger network response elicited on average by old-relative to new-in-context items (Fig. 1F, differences between averages in firing rates induced by pairs of old- vs. new-in-context items, Δf_old-new,_ are positive, and hence old items elicited stronger response, see Pairwise differences section in Methods). To gain further insights in ways inaccessible to in-vivo experiments, we examined the synaptic strength of the within- and between-network connectivity, and neuronal excitability dynamics (BCPNN bias, see Methods). The high odor recall performance cannot be explained by observations in the within-network connectivity, as the differences in average within-network connectivity between pairs of old vs. new items, Δw_old-new_, drifts towards negative values (Fig. 1G, see Pairwise differences section in Methods), primarily due to high recency of new items which boosts their connectivity. Regarding the bias factor, old-in-context items were typically presented more times than new-in-context items, as a consequence of task design, the cell assemblies corresponding to more repetitive old-in-context items exhibited higher neuronal excitability. The distribution of the differences in average bias between pairs of old vs. new items, Δb_old-new_, is positive, and favors old items in Fig. 1H, which partly contributed to the odor recall performance. Prior to the Memory Assessment block, items that were presented in context A established an excitatory associative binding (Fig. 1I, top right, EPSPs in middle) unlike other items, never cued in context A beforehand, were subjected to disynaptic inhibition (Fig. 1I, top left, IPSPs in bottom, see Methods). During the Memory Assessment block, which was still part of the continual learning process in context A, items that were initially new to that context became old-in-context items once they were used as a stimulus in the Memory Assessment. Hence, plastic disynaptic inhibition built during the earlier Item-Context Encoding block was transformed to excitatory binding (continual learning process) after the odor item was cued in the Memory Assessment block (see Methods). All in all, the synaptic weights of the associative item-context binding and the bias factor contributed to the observed difference in firing rates between old vs new items in Figure 1F.

Next we challenged our model by simulating the extended task with three context transitions (A ➞ B ➞ A ➞ Memory Assessment block in context B), as proposed by Panoz-Brown et al. (2016). In their second experiment (denoted as Experiment 2), 8 out of 16 odors were stimulated in context A (as before in Experiment 1), followed by 8 odors in context B. After transitioning to context A again, the remaining 8 items not shown in context A yet were presented. The Memory Assessment part of Experiment 2 was then conducted in context B, unlike in Experiment 1, with new-in-context items presented along with the previously encoded old-in-context items, as before (Fig. S1, an example of a three-context-transition task). The same model, i.e. without any further re-tuning, reproduced again quantitatively similar odor recall performance as in Panoz-Brown et al.’s (2016) Experiment 2 (Fig. 1E; model data: mean=82.35, SD=3.49, n=119, mean represents the total number of successes across all n-trials, SD derived from the Bernoulli distributions for the probabilities of successes across all n-trials, n corresponds to simulated old-new pairs, and experimental data: mean≈84, SD≈5.3, mean reflects the average performance of rats in 9 sessions, combining the initial and terminal sessions, and SD reflects the average standard error of the mean (SEM) across rats for the combined initial and terminal sessions).

### Synergy of episodic memory and recency: An alternative task design (Arrangement 2)

To prevent rats from utilizing any semantic rules concerned with items’ recency, Panoz-Brown et al. (2016) randomly intermingled Arrangement 1 trial blocks with trials of an alternative structure called Arrangement 2. In Arrangement 2, the order of the item presentation in the Item-Context Encoding block prior to the Memory Assessment was switched so that old-in-context items featured higher recency than the new-in-context items (Fig. 2A, Arrangement 1 vs Arrangement 2). Panoz-Brown et al. (2016) did not report any experimental results for Arrangement as recency effects might be confounded with context dependent episodic memory. We nevertheless wanted to provide a qualitative prediction about the memory performance in the Arrangement 2 task given the synergy of short-term recency and episodic memory.

**Figure 2:**
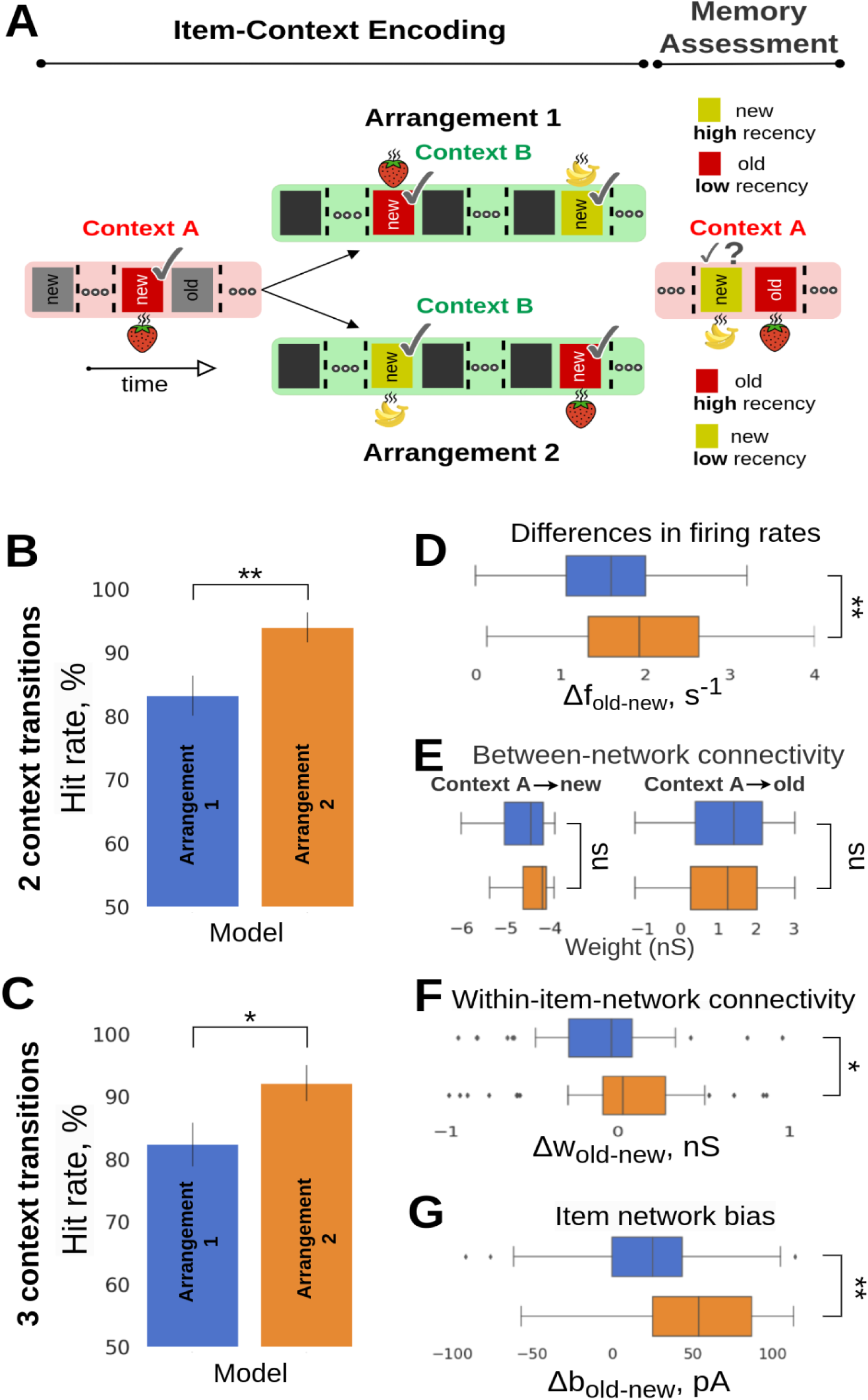
**A**, Structure of the episodic memory task: Arrangement 1 vs. Arrangement 2. In Arrangement 1 (cf. Fig. 1B), the new-in-context item was encoded more recently than the old-in-context item prior to the Memory Assessment phase. This order was reversed in Arrangement 2. **B,C** Model performance in new- vs. old-in-context memory discrimination between Arrangements 1 and 2 for two- and three-context transitions, respectively. Error bars represent SDs derived from the Bernoulli distributions for the probabilities of success (hit) across all trials (scaled to %). **D**, Boxplot of the differences in average firing rates between pairs of old vs. new items, Δf_old-new_, in Arrangement 2 (bottom) and in Arrangement 1 (top). The trial-average firing rates represent the means of evoked spiking frequency during the activation of a given item during a test trial in the Memory Assessment phase. **E**, Synaptic connectivity (AMPA and slower NMDA receptor mediated weights) between Item and Context networks is similar in both arrangements and remains resistant to changes (i.e., order of item activation) due to encoding with long-term Hebbian plasticity (τ_p_=30 s, Table 1). **F**, Distribution of the differences in average within-network connectivity between pairs of old vs. new items, Δw_old-new_, for the Arrangement 1 (blue) and Arrangement 2 (orange). **G**, Distribution of the differences in average intrinsic excitability between pairs of old vs. new items, Δbias_old-new_, for the Arrangement 1 (blue) and Arrangement 2 (orange). Both within-network connectivity and bias effects of old items within the Item network are stronger in Arrangement 2 due to recency, thus leading to higher performance as observed in B.

The model predicted indeed higher odor recall performance in Arrangement 2 than in Arrangement 1 for both two- (Fig. 2B, Arrangement 1: mean=83.21, SD=3.12, n=143 vs Arrangement 2: mean=93.33, SD=2.39, n=99; p<0.01, Fisher’s exact test), and three-context transitions (Fig. 2C, Arrangement 1: mean=82.35, SD=3.49, n=119 vs Arrangement 2: mean=92.13, SD=2.85, n=89; p<0.05, Fisher’s exact test). To elucidate the synaptic origins and network correlates of the performance enhancement, we analyzed key model variables such as spiking activity of excitatory units representing the old- and new-in-context items, synaptic strength of the within- and between-network connectivity, and neuronal excitability dynamics (BCPNN bias, see Methods). We observed that the differences between the averages in firing rates induced by old- vs. new-in-context items, Δf_old-new,_ increased significantly in Arrangement 2 relative Arrangement 1 (Fig. 2D; p<0.01, two-sample t-test, 90 simulated trials [pairs of odors in the Memory Assessment] in Arrangement 1 and 62 in Arrangement 2, see Pairwise differences section in Methods), implying the improved capability of the model to discriminate and accurately identify new-in-context odor items. We partially attributed this to the temporary enhancement in the strength of the within-network connectivity (Fig. 2F, p<0.05, Mann–Whitney U test, 47 simulated trials [pairs] in Arrangement 1 and 41 trials in Arrangement 2). As mentioned earlier, the within-network connectivity was preloaded (long-term memory representations of items and contexts were encoded prior to the Item-Context Encoding block), so it was short-term synaptic augmentation that rapidly upregulated the effective synaptic weights. This enhancement was short-lasting, limited by the augmentation time constant, and thus it could only be effective when the stimulation of a given item in context B was within a narrow time window relative to the temporal scales of the Memory Assessment block in context A. Furthermore, we observed a notable increase in the difference between the neuronal excitability (Δbias) for old- vs. new-in-context items in Arrangement 2 (Fig. 2G, p<0.01, Mann–Whitney U, 47 simulated trials [pairs] in Arrangement 1 and 41 trials in Arrangement 2). Old-in-context items were stimulated more often than the new ones as a result of the altered task structure (Fig. 2A). In Arrangement 2 the final stimulation of an old-in-context item had to take place after the most recent activation of a new-in-context item even if there were cases that old-in-context items had been activated before (i.e., strawberry [old, first stimulation] – banana [new, first stimulation] – strawberry [old, second stimulation]). Therefore, by enforcing Arrangement 2 the intrinsic neural excitability dynamics of old items was enhanced. It is worth mentioning that modification of the temporal order of items between Arrangement 1 and Arrangement 2 did not yield any meaningful change for the between-network connectivity (Fig. 2E, p>0.05, Mann–Whitney U, 47 simulated trials [pairs] in Arrangement 1 and 41 trials in Arrangement 2). The between-network connectivity was long lasting (*τ*_p_=30 s, Table 1) and resistant to small temporal changes to support episodic retrieval. In general, we highlight the importance of these long-lasting synaptic traces to perform item-context association tasks, as we observed that fast Hebbian plasticity with short time synaptic constants (e.g., a time constant *τ*_p_ of 5 s was commonly employed in working memory settings by other models) alone could not solve the task, as the temporal memory traces of previously encoded item-context pairs decayed rapidly.

### Unbalanced training paradigm with two context transitions

To further exploit the predictive capabilities of the model, we examined the effect of the frequency of stimulus presentation (multiplicity or repetition of stimuli) as a potential factor modulating the item familiarity on the item-in-context recall. In particular, we set out to study if the stimulus multiplicity on top of the recency would synergistically outcompete the episodic memory effect in the old- vs new-in-context item choice, thereby leading to the higher recall error rate. To this end, we resorted to Arrangement 1 (competition between recency and episodic memory phenomena). However, unlike in the balanced Experiment 1, we now increased the number of new-in-context odor presentations resulting in an unbalanced scenario where new items were presented more frequently (i.e., in Fig. 3A in the Memory Assessment block, the new-in-context-A item (yellow) had been used twice in the preceding context B, while the old-in-context-A item (blue) had appeared only once). Surprisingly, our expectation that the enhanced familiarity due to increased multiplicity along with recency should outcompete episodic memory and “mislead” the model in the old- vs new-in-context choice during the Memory Assessment turned out to be false. In fact, we found evidence of comparably high performance for the unbalanced training task, i.e. reference task: mean=83.21, SD=3.12, n=143 vs unbalanced training task: mean=83.92, SD=2.83, n=168; p>0.05, Fisher’s exact test.

**Figure 3:**
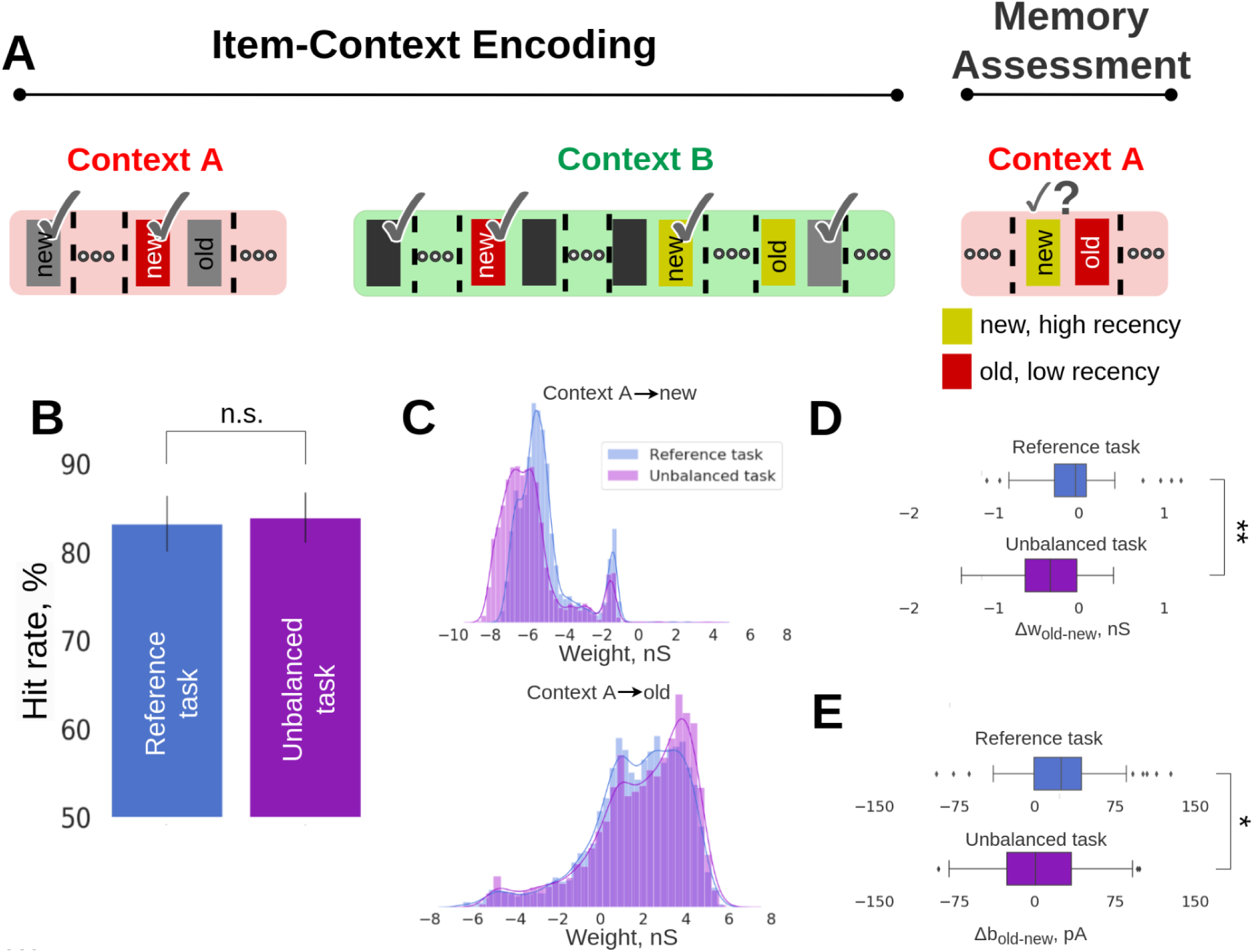
Unbalanced training prediction task. A, Schematic of the unbalanced training task. As in the reference two-context-transition task (Fig. 1B), half of the items were presented in context A, followed by the presentation of all the items in context B. The Memory Assessment was conducted in context A by presenting pairs of new-old items. However, for the unbalanced training task, we stimulated more times the new-in-context-B items than in the corresponding reference task. B, Average recall performance (hit rate, %) for the reference and unbalanced prediction task corresponding to Arrangement 1 configuration. SDs derived from the Bernoulli distributions for the probabilities of success (hit) across all trials (scaled to %).C, Distributions of associative weights (AMPA and slower NMDA receptor mediated weights, reported prior to the Memory Assessment phase) from context A to new-in-context-A items (top, disynaptic inhibitory weights), and from context A to old-in-context-A items (bottom) for the reference and unbalanced training prediction task. D, Boxplot of the differences in average within-network connectivity between old vs. new items, Δw_old-new_, for the reference task with two context transitions (top) and the unbalanced training scenario (bottom). E, Boxplot of the differences in average intrinsic excitability (neuronal bias) between pairs of old vs. new items, Δb_old-new_, for the reference task with two context transitions and the unbalanced training scenario.

We next sought to mechanistically explain the comparable performance in the unbalanced task, which was opposite to our expectation as we predicted lower performance. First, we analyzed the between-network connectivity and found that disynaptic inhibition between context A and the new-in-context items was strengthened (Fig. 3C, top, p=1.9 x 10^-268^, Mann-Whitney U-test, 6638 weights from Context A to all the new items for the reference task vs. 5382 weights from Context A to all the new items for the unbalanced task). This effectively resulted in a more negative (inhibitory) association in the unbalanced training task. There were more opportunities for new-in-context items to be repeated in context B of the Item-Context Encoding block than in the reference task setup. This strengthened not only their associative binding with context B but also their dissociation with context A (mediated by plastic disynaptic inhibition, see Methods). At the same time, associative excitatory binding between context A and those items that were later considered old-in-context in the Memory Assessment block became stronger, predominantly due to their less frequent presentation as a stimulus in the competing context B. Indeed, in the spirit of Bayesian nature of BCPNN learning, a more specific and exclusive pairing of two memory patterns induces stronger associative binding. The observed changes of the between-network connectivity (Fig. 3C, bottom, p=7.82 x 10^-16^, Mann-Whitney U-test, 6725 weights from Context A to all the old items for the reference task vs. 5464 weights from Context A to all the old items for the unbalanced task) can explain the reason why the hit rates were not reduced for the unbalanced training task (Fig. 3B). However, differences in the within-network connectivity and bias still hurt recall, because new items-in-context featured stronger attractor connectivity (Fig. 3D, enhanced within-network connectivity of new items in the unbalanced task drove the Δw_old-new_ distribution towards more negative values, reference vs. unbalanced task: p<0.01, Mann-Whitney U-test, n_Reference_=47, n_Unbalanced_=59, see Pairwise differences section in Methods), and boosted neuronal excitability (bias, Fig. 3E, references vs. unbalanced task: p<0.05, Mann-Whitney U-test, n_Reference_=47, n_Unbalanced_=59) compared to the original task design due to the additional stimulus repetitions. These effects led to a stronger competition between non-episodic and episodic memory effects compared to the reference task. Still, our simulations showed that episodic memory processes reflected in associative between-network binding overpowered familiarity and recency effects manifested at the item representation level.

Collectively, the alterations in context-item binding, reflected in the between-network connectivity weights and caused by varying multiplicity of item presentations, resulted in maintaining a comparable high item-in-context memory performance. Due to the important role of the aforementioned disynaptic inhibition between context A and new-in-context items, we conducted a follow-up experimental manipulation by severing the disynaptic weights connecting the networks for the unbalanced task. Subsequent recall rates in the Memory Assessment block showed a dramatic decrease in performance (Fig. S2, unbalanced task: mean=83.92, SD=2.83, n=168 vs “No disynaptic inhibition” task: mean=16.66, SD=5.37, n=48; p<0.001, Fisher’s exact test). Similar manipulation (e.g., removal of between-network disynaptic inhibitory weights) to the reference task leads to poor performance as well.

### Memory interference by an additional episodic context

From the behavioral perspective on episodic memory it is interesting to study the effect of memory interference by introducing yet another context C (arena) just preceding the Memory Assessment block (Fig. 4A). Increasing the complexity of the task (extended memory and temporal demands) by introducing additional contexts is a method often used in item-in-context episodic memory tasks on rats, and this process typically leads to lower performance scores (Weisz et al., 2012). Our intention was to make behaviourally relevant predictions about the odor recall performance in a more challenging setup compared to the reference task, and quantify potential behavioral changes in performance. In line with previous related behavioral experiments, the recall performance in our extra context task dropped significantly compared to the reference task (Fig. 4B, reference task: mean=83.21, SD=3.12, n=143 vs extra context task: mean=70.7, SD=5.02, n=82; p<0.05, Fisher’s exact test).

**Figure 4:**
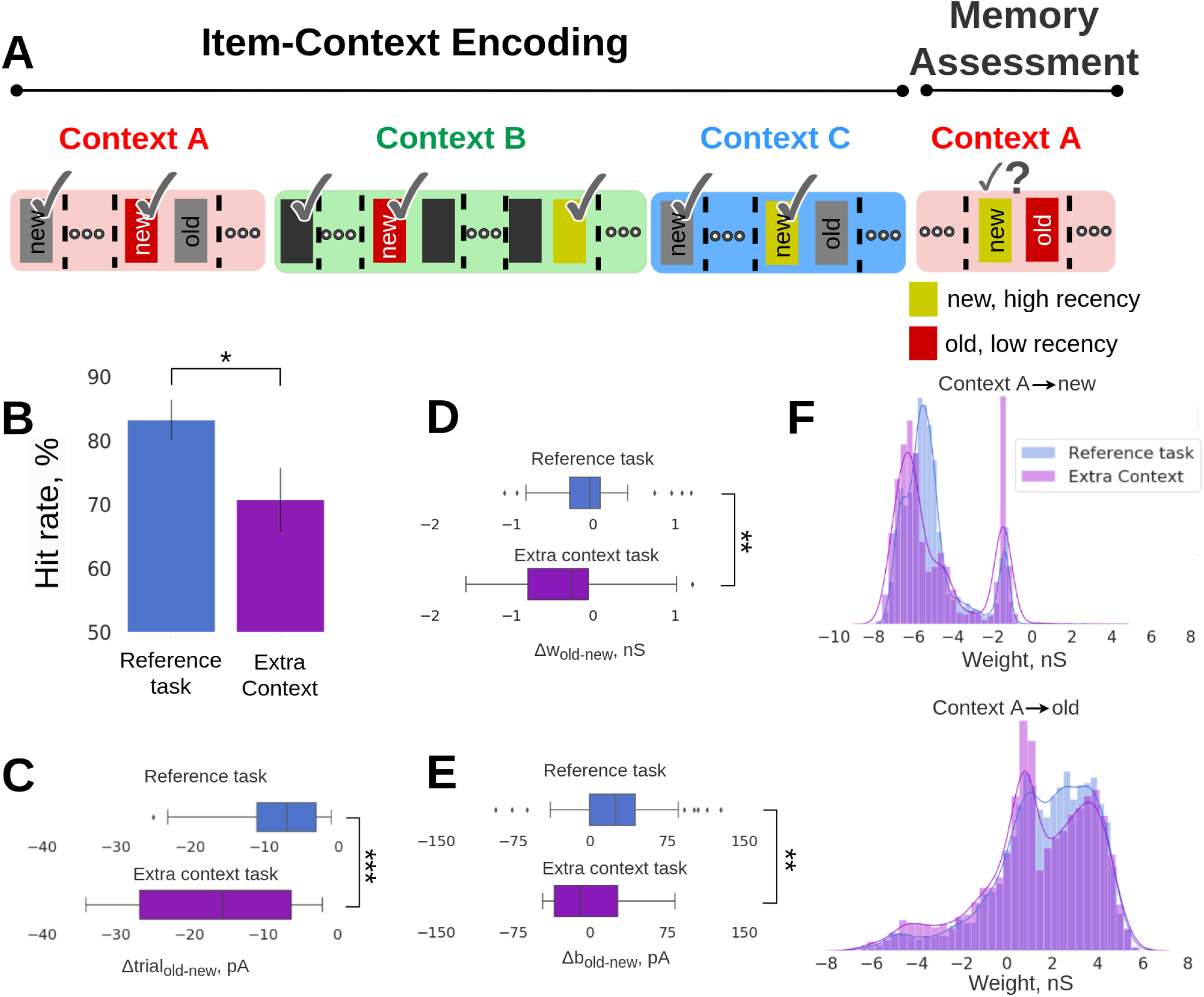
Extra context prediction task. A, Schematic of the extra context task. Introducing an extra context (blue) in the Item-Coding Encoding block prior to the Memory Assessment block, in which we simulated randomly half of the 16 items in new-old item pairs. B, Average recall performance (hit rate, %) for the reference and extra context prediction tasks corresponding to Arrangement 1 configuration. SDs derived from the Bernoulli distributions for the probabilities of success (hit) across all trials (scaled to %). C, Boxplot of the differences in average trial-index between pairs of old vs. new items, Δtrial_old-new_, for the reference task with two context transitions (top) and the extra context task (bottom). As a trail-index we define the trial index of the most recent activation of the items (i.e., a new item with trial-index=40 is more recently encoded compared to an old item with trial-index=30, and their relative recency difference is Δtrial_old-new_=-10). When an item was activated at the very first trial block, it was assigned with a trial-index=1, and once it was activated again at a later trial, the trial-index was flexibly updated to correspond to the last activation position. The figure shows that the difference in trial-index between old and new items becomes more negative for the extra context task, thus increasing their relative recency. D, Boxplot of the differences in average within-network connectivity between pairs of old vs. new items, Δw_old-new_, for the reference task with two context transitions (top) and the extra context task (bottom). E, Boxplot of the differences in average intrinsic excitability (neuronal bias) between pairs of old vs. new items, Δbias_old-new_, for the reference task with two context transitions and the extra context scenario. F, Distributions of associative weights (reported prior to the Memory Assessment block) between context A and new-in-context-A items (top, disynaptic inhibitory weights), and context A and old-in-context-A items (bottom) for the reference and extra context tasks.

The new simulated task puts short-term recency effects in conflict with episodic memory, just as in the behavioral task with Arrangement 1, though in a more complex and longer item-in-context configuration facilitated by an extra context C. In particular for this new context C, we cued randomly 8 of the available 16 memory items (see Fig. 4A) following an analogous procedure of presenting items as in Experiment 1 (i.e., Fig. 1C, 0-15 s). Later, items that had been cued in context C, could be used in the Memory Assessment block. Given the Arrangement 1 criteria (“new-in-context items should be more recently encoded than the old items”), we observed that new-in-context-A items (from the perspective of Memory Assessment) were the items that were mainly activated in the extra context C (latest presentation before the Memory Assessment block) as opposed to the old-in-context-A items whose most recent presentation took place in context B (not in context C, Fig. 4A). This was an emerging outcome of the new task setup combined with Arrangement 1 requirements. Since old items were rather infrequent in trials belonging to context C as opposed to new items that were activated extra times in context C, there was a longer temporal distance between the trials of the most recent activation of a new and its old item pair (higher relative recency between pairs of items, Fig. 4C, reference vs. extra context task: p<0.001, Mann–Whitney U test, n_Reference_=47, n_ExtraContext_=34). The longer emerging temporal distance (higher relative recency) between old and new items, led to enhanced within-network connectivity for new-in-context items (Fig. 4D, Δw_old-new_ distribution drifts to more negative values indicating within-network connectivity enhancement of new items, reference vs. extra context task: p<0.01, Mann-Whitney U-test, n_Reference_=47, n_ExtraContext_=34, see Pairwise differences section in Methods). Also, the extra activations of new-in-context items in context C yielded stronger learned intrinsic excitability (higher multiplicity hypothesized to enhance familiarity in experimental memory settings) compared to the reference task (Fig. 4E, Δb_old-new_ distribution drifts to negative values, reference vs. extra context task: p<0.01, Mann-Whitney U-test, n_Reference_=47, n_ExtraContext_=34). The above changes in within-network connectivity and intrinsic excitability boosted spiking activity of new items making it harder for the model to distinguish between new- and old-in-context items (selection was made based on significant spiking activities differences between old and new items), and hence these synaptic- and neuronal-level changes may explain the observed performance decline. Last but not least, we observed a similar disynaptic inhibition trend for the extra context task as in the previous unbalanced training task (Fig. 3C, top), that is, stronger disynaptic inhibition from context A to all the new-in-context-A items compared to the reference task (Fig. 4F, top, reference vs. extra context task: p=5.32 x 10^-8^, Mann-Whitney U-test, n_Reference_=6638 weights, n_ExtraContext_=5402 weights). However, the between-network connectivity (associative episodic item-in-context binding) was weaker compared to the reference task. Even though old items were activated infrequently in the extra context C, still they were activated more times in other contexts compared to the reference task, and hence this additional repetition in other contexts can weaken the associative binding as shown in Chrysanthidis et al. (2022) study (Fig. 4F, bottom, reference vs. extra context task: p=4.7 x 10^-13^, Mann-Whitney U-test, n_Reference_=6725 weights from context A to all the old items, n_ExtraContext_=5453 weights from context A to all the old items). It is worth noting that the majority of old items (in context A) that were cued in context C followed the stimulation logic of Arrangement 2 (Fig. 2A), i.e. old-in-context items were more recently encoded than the new ones, and were excluded from the Arrangement 1 analysis.

### Reverse training task

In the original reference task (Fig. 1B) new-in-context items were rewarded only once upon selection since after the reward they were treated as old items, and no further reward was provided even after another presentation in the same context. Therefore, the overall reward was distributed uniformly across odors in both contexts in the Item-Context Encoding prior to the Memory Assessment block. By reversing the reward scheme and utilizing a rule to provide rewards only to old-in-context items, we can introduce a reward imbalance between items as an old item can be rewarded as many times it is activated in the context (Fig. 5A).

**Figure 5:**
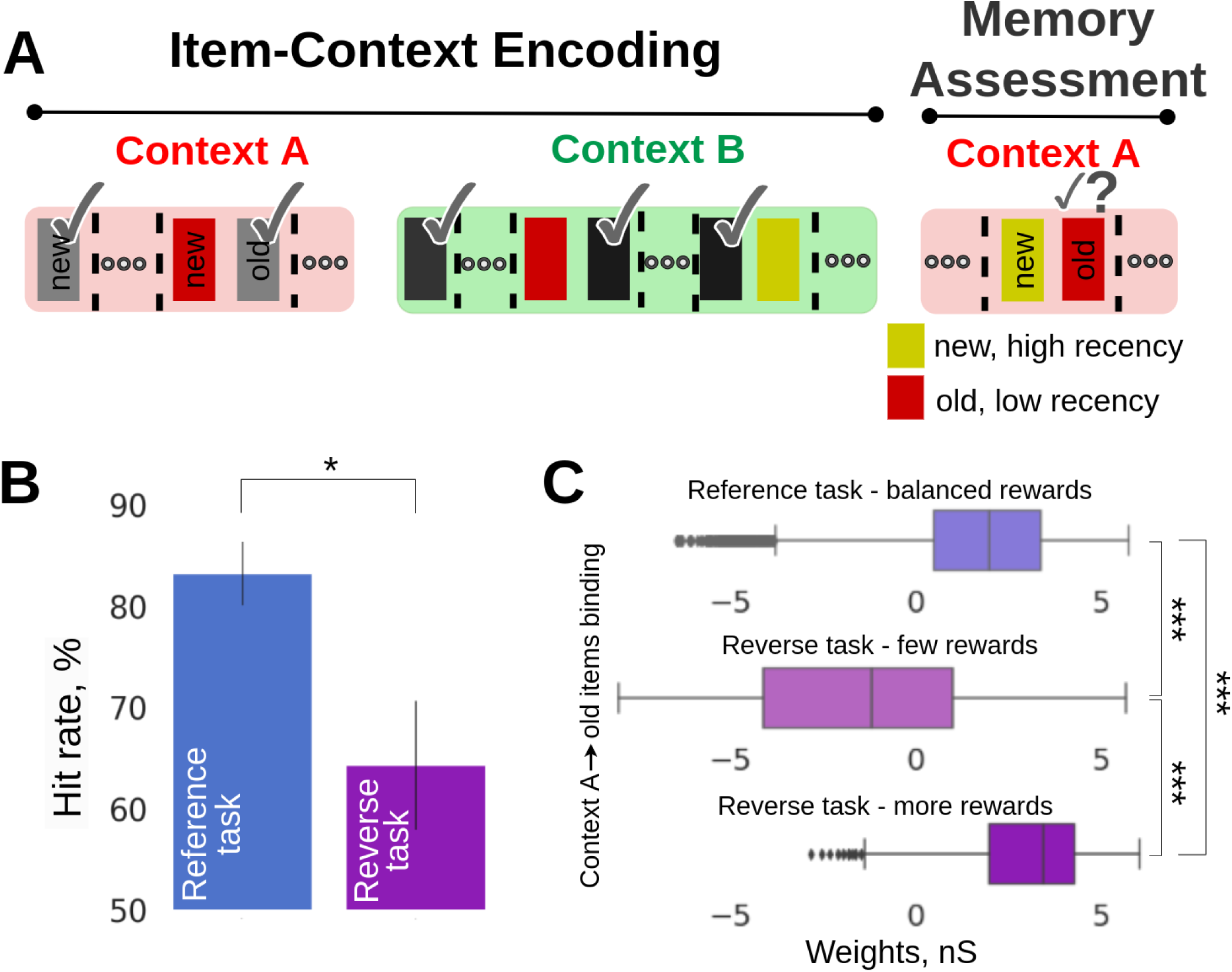
Reverse training prediction task. **A**, Schematic of the reverse training task. which is similar to the reference task with the only difference that old items-in-context are rewarded instead of new-in-context items. **B**, Average recall performance for the suite of the prediction task. Data is shown for Arrangement 1. SDs derived from the Bernoulli distributions for the probabilities of success (hit) across all trials (scaled to %). **C**, Boxplots of the differences in the between-network connectivity (associative weights, AMPA and slower NMDA receptor mediated weights, reported prior to the Memory Assessment block) between context A and old-in-context-A items for the reference task (top), between context A and old-in-context-A items that were rewarded few times during the reverse training task (middle), and between context A and old-in-context-A items that were rewarded multiple times during the reverse training task (bottom). Old-in-context-A items that were rewarded multiple times (i.e., more than two rewards) in the reverse training task featured stronger synaptic connectivity with context A (bottom) than the corresponding ones rewarded fewer times in the reverse training task (middle).

The reverse training task substantially hurt model performance (Fig. 5B, reference task: mean=83.21, SD=3.12, n=143 vs. reverse training task: mean=64.28, SD=6.4, n=56; p<0.05, Fisher’s exact test). The poor memory performance is best explained by the between-network connectivity, which affects item-in-context memory performance, as observed in earlier prediction tasks. The reward imbalance resulted in varying levels of between-network connectivity strength among old-in-context items, promoting less robust associative episodic memory binding for some old items (Fig. 5C, difference between the means of associative weights between reference task and reverse task; cases with more and few odor rewards, i.e. reference: reward balance, vs reverse training task: reward imbalance; more rewards, p<0.001, Mann–Whitney U test, n_Reference_=6725, n_more-Rewards_=2154, n represents the weights from context A to all the old items; reference: reward balance, vs reverse training task: reward imbalance; few rewards, p<0.001, Mann–Whitney U test, n_Reference_=6725, n_few-Rewards_=712; and reverse training task: reward imbalance; more rewards, vs reverse training task: reward imbalance; few rewards, p<0.001, Mann– Whitney U test, n_more-Rewards_=2154, n_few-Rewards_=712.

Weak between-network connectivity for old-in-context-A items that were rewarded fewer times decoupled the Item-Context networks (Fig. 5C). On the other hand, when old items were rewarded multiple times, their between-network connectivity was strengthened at the cost of other coupled items (in context A), because learning was continuous throughout the task. Bayesian learning normalizes and updates weights continuously over estimated presynaptic (Bayesian-prior) as well as postsynaptic (Bayesian-posterior) spiking activity. We noticed that the reward imbalance in the reverse training task, particularly for these cases where old items were rewarded less often led to incorrect odor choices.

## Discussion

Testing episodic memory is important to elucidate the mechanisms that could interfere with, enhance or impact this memory system. While there is a wealth of research on the behavioral manifestations of this type of memory (Wilson et al., 2013; Lesburguères et al., 2017; Kanatsou et al., 2016), especially in rats using item-in-context paradigms, little is known about the underlying neural mechanisms that govern their interactions, and how these learning effects with different temporal characteristics interplay at a network level. In a pivotal experimental task on episodic memory by Panoz-Brown et al. (2016), rats demonstrated high accuracy in solving an item-in-context task, indicating their reliance on episodic memory. Motivated by this item-in-context episodic task, we constructed a computational spiking neural network model to explore the neural mechanisms that govern the intricate interplay between episodic and short-term recency memory effects at a mesoscopic network level. We attributed Panoz-Brown et al.’s (2016) behavioral findings to emergent network dynamics resulting from local synaptic plasticity phenomena operating across various timescales. Our objective was to offer mechanistic insights into these computationally underexplored synergistic memory phenomena. It should be noted that in our computational study we deliberately and consistently referred to the short-term memory phenomena of interest as recency rather than familiarity, used originally by Panoz-Brown et al. (2016). We consider recency as a more precise term than familiarity even if the latter has been linked to the general notion of memory strength affected among others by stimulus recency (Yonelinas et al., 2010).

### Model predictions and experimental data

Our dual network model successfully matched empirical observations of item-in-context memory in rats (Fig. 1D, E), as reported by Panoz-Brown et al. (2016). Notably, this was achieved while maintaining biologically constrained network connectivity, postsynaptic potential amplitudes, and firing rates compatible with mesoscale recordings from cortex and earlier models. We also generated predictions regarding behavioral outcomes in three modified task paradigms, which could be examined in a follow-up experimental study. In particular, we sought to explore a wider scope of recency effects in episodic memory retrieval.

In our first simulated prediction task we switched the order of odor presentation such that old items-in-context were more recently encoded than new items. We then quantified the increase in memory recall performance due to the synergistic contribution of episodic and short-term memory effects of recency. Our findings align with similar experiments in rats that focus on the potential impact of recency on the Object-in-Context (OIC) task (Tam et al., 2015). In that experimental study rats first freely explored an object-i in a visual context-X, and then explored another object-j in context-Y. During a subsequent memory test phase the rats were supposed to choose between object-i or object-j either in context-X or context-Y. The data revealed evidence of enhanced performance when the two items were tested in context-Y (compared to context-X). A contributing factor could be that object-j was more recently encoded than object-i, resulting in a relatively stronger memory trace for object-j compared to object-i at the time of the test. Their reasoning aligns with our mechanistic explanation as we observed strengthened memory traces for recently encoded items (see Fig. 2F).

Our third prediction task with an additional third context could be related to Weisz et al.’s (2012) experiments aimed at examining the impact of memory and time overload on the capacity to recognize new-in-context items. In their experimental protocol rats first freely explored in an open field two non-identical items in different visual contexts labeled as A, B, and C for 5 minutes each. The subsequent test session took place in one of the arenas (A, B, or C) with two copies of the same object, where only one matched the original spatial location. To receive a reward rats had to identify the new combination of context-object-place with the specific arrangement of the objects. The rats were first tested using two contexts (A, B) and later, in a separate trial, an additional context C was introduced (A, B, C) expanding the complexity of the task. The data shows higher recognition of new-in-context objects for the two context scenario (A, B) compared to the three context scenario (A, B, C) evidencing less recall with increasing requirements. We saw a similar decrease in memory performance in our simulation task, when we increased the number of contexts from two to three.

### BCPNN vs. STDP discussion

Our model uses the Bayesian-Hebbian associative learning rule (BCPNN) while there are other associative Hebbian-like learning rules more commonly used in computational studies, e.g. spike-timing dependent plasticity (STDP) (Ren et al., 2010; Rossum et al., 2000). We cannot exclude that an alternative learning rule like STDP may, in principle, mechanistically explain item-in-context memory. However, the key strength of our BCPNN rule lies in the intrinsic regulation of spiking activity through synaptic learning of long-lasting disynaptic inhibition (via double bouquet cells which may play an important role in shaping neural activity and circuitry, DeFelipe et al., 2006; Krimer et al., 2005; Kelsom and Lu, 2013; Chrysanthidis et al., 2019), and contrasts with known issues of network stability and robustness with STDP. Indeed, in the absence of disynaptic inhibition our network fails to solve the task as a result of emerging instabilities (Fig. 3B). STDP predominantly operates on the millisecond scale and even if synapses were depotentiated they would not represent any meaningful learned long-term disynaptic inhibitory component. We do not exclude that a similar model relying on STDP-tuned connectivity that also includes disynaptic inhibition could perform well. However, in the classical STDP models disynaptic inhibition is rarely integrated.

### Related Models of episodic memory

Various computational models, notably dual-process models, have investigated the processes of familiarity and recollection (Wixted, 2007), sometimes within the framework of a single memory trace (Greve et al., 2009). Our research diverges from conceptualizing recency solely as a familiarity process to examine the impact of short-term dynamics on recall. On the whole, computational models of episodic memory remain relatively scarce (Norman and O’Reilly, 2003, Brea et al., 2023), often integrating abstract or non-spiking representations. Furthermore, the subset of models specifically addressing the interplay between short-term dynamics and episodic memory is even more limited. Notably, a recent model showed that selectively encoding episodic memories at the end of an event led to better subsequent prediction performance (Lu et al., 2022). Furthermore, in more dated investigations, temporal context models in the domain of episodic memory have demonstrated a broad spectrum of recall phenomena including recency and contiguity effects observed across immediate, delayed, and continuous distractor-free recall scenarios (Sederberg et al., 2008).

### Conclusion

One key strength of computational modeling is that it can bridge spatial scales, from behavior and whole-brain dynamics to single-cell activity and thus explain more data. Our detailed spiking model bridges these perspectives and represents a novel computational attempt to connect neural and synaptic processes with mesoscopic manifestations underpinning complex effects of short-term memory dynamics on episodic memory recall, and item-in-context memory, in particular. We have shown a quantitative match with Panoz-Brown et al.’s (2016) experimental findings obtained in a detailed spiking network model, constrained by available biological data (constrained network connectivity with neurobiologically plausible postsynaptic potentials, firing rates, and other parameters). We consider this to be a significant step towards bridging the gap between behavioral correlates of complex episodic memory phenomena and the underlying synaptic mechanisms.

## Methods

### Neuron and synapse model

We use adaptive exponential integrate-and-fire point model neurons, which feature spike frequency adaptation, enriching neural dynamics and spike patterns, especially for the pyramidal cells (Brette and Gerstner, 2005). This neuron model is an effective model of cortical neuronal activity, reproducing a wide variety of electrophysiological properties, and offers a good phenomenological description of typical neural firing behavior, but it is limited in predicting the precise time course of the subthreshold membrane voltage during and after a spike or the underlying biophysical causes of electrical activity (Gerstner and Naud, 2009). We slightly modified it for compatibility with the BCPNN synapse model (Tully et al., 2014) by integrating an intrinsic excitability current.

Development of the membrane potential *V_m_* and the adaptation current *I_w_* is described by the following equations:

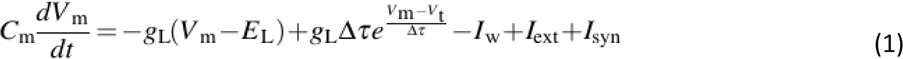

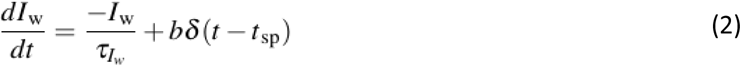

Equation 1 describes the dynamics of the membrane potential *V_m_* including an exponential voltage dependent activation term. A leak current is driven by the leak reversal potential *E_L_* through the conductance *g_L_* over the neural surface with a capacity *C_m_*. Additionally, *V_t_* is the spiking threshold, and Δ_T_ shapes the spike slope factor. After spike generation, membrane potential is reset to *V_r_*. Spike emission upregulates the adaptation current by *b*, which recovers with time constant *τ_Iw_* (Table 1). To simplify the model, we have removed subthreshold adaptation, which is part of some AdEx models.

Besides a specific external input current *I_ext_*, model neurons receive synaptic currents *I_synj_* from conductance based glutamatergic and GABA-ergic synapses. Glutamatergic synapses feature both AMPA/NMDA receptor gated channels with fast and slow conductance decay dynamics, respectively. Current contributions for synapses are described as follows:

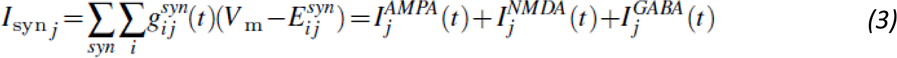

The glutamatergic synapses are also subject to synaptic depression and augmentation with a decay factor *τ_D_* and *τ_A_*, respectively (Table 1), following the Tsodyks-Markram formalism (Tsodyks and Markram, 1997). We have chosen those time-constants from the plausible range of computational fits made on the basis of electrophysiological recordings of cortical pyramidal cells (Wang et al., 2006). The utilization factor u represents the fraction of available resources used up by each transmitted spike (a proxy of synaptic release probability), whereas x tracks the fraction of resources that remain available due to transmitter depletion (synaptic depression):

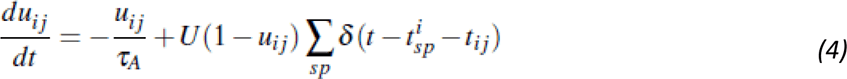

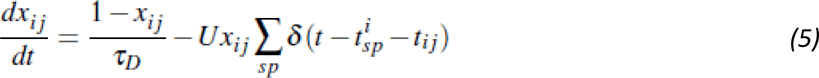

### Spike-based BCPNN plasticity

We implement synaptic plasticity of glutamatergic synapses using the BCPNN learning rule (Lansner and Ekeberg, 1989; Wahlgren and Lansner, 2001; Tully et al., 2014). BCPNN is derived from Bayes rule, assuming a postsynaptic neuron employs some form of probabilistic inference to decide whether to emit a spike or not. Despite that it accounts for the basic Bayesian inference, it is considered more complex than the standard STDP learning rule (Caporale and Dan, 2008), and as such it reproduces the main features of STDP plasticity. In a previous study, we demonstrated that with BCPNN synaptic plasticity, but not with standard Hebbian STDP, the model can reproduce traces of semantization as a result of learning (Chrysanthidis et al., 2022). Therefore, in our effort to explore the interplay of episodic memory with recency effects we utilize the BCPNN learning rule.

The BCPNN synapse continuously updates three synaptic biophysically plausible local memory traces, *P_i_*, *P_j_* and *P_ij_*, implemented as exponentially moving averages (EMAs) of pre-, post- and co-activation, from which the Bayesian bias and weights are calculated. EMAs prioritize recent patterns, so that newly learned patterns gradually replace old memories. Specifically, learning implements exponential filters, Z, E, and P, of spiking activity with a hierarchy of time constants, *τ*_Z_, *τ*_e_, and *τ*_p_, respectively. Due to their temporal integrative nature they are referred to as synaptic (local memory) traces.

To begin with, BCPNN receives a binary sequence of pre- and postsynaptic spiking events (*S_i_*, *S_j_*) to calculate the traces *Z_i_* and *Z_j_*:

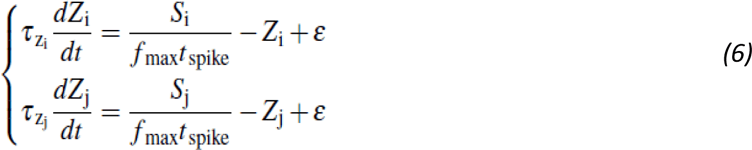

*f_max_* denotes the maximal neuronal spike rate, *ε* is the lowest attainable probability estimate, *t_spike_* denotes the spike duration while *τ*_Zi=_*τ*_Zj_ are the presynaptic and postsynaptic time constants, respectively (*τ*_Z_ =*τ*^AMPA^ =5 ms for AMPA, and *τ*_Z_ =*τ*^NMDA^ =100 ms for NMDA components, Table 1).

*E and P* traces are then estimated from the *Z* traces as follows:

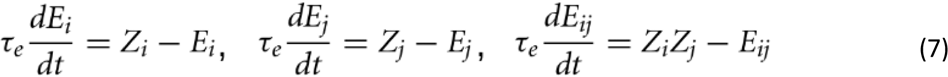

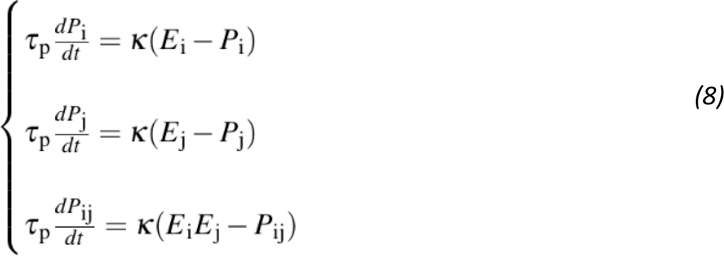

The parameter *κ* adjusts the learning rate, reflecting the action of endogenous modulators of learning efficacy (i.e., activation of a D1R-like receptor). Setting *κ*=0 freezes the network’s weights and biases, though in our simulations the learning rate remains constant (*κ*_*normal*_=0.3) during encoding. However, to account for the experimental paradigm we trigger a transient increase of plasticity to simulate the impact of a reward signal on the memory system by implementing eligibility traces (see Eq. 7) and upregulating the associative plasticity gain (κ_reward_=1) upon successful execution of the task by the model.

Finally, *P_i_*, *P_j_* and *P_ij_* are used to calculate intrinsic excitability *β_j_* and synaptic weights *w_ij_* with a scaling factor *β_gain_* and 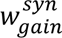 respectively (Table 1):

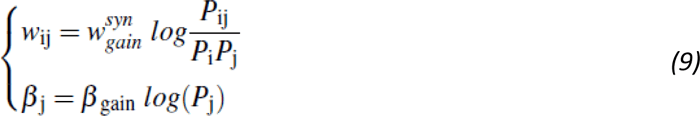

BCPNN is a Hebbian-like learning rule, neurons that are coactive are coupled with excitatory connectivity. However, neurons that do not fire together in a certain time window feature low coactivation traces (P_ij_), and based on Equation 9, the final weight update will produce negative conductance. The negative binding is interpreted as disynaptic inhibition mediated by dendritic targeting regular spiking non-pyramidal (RSNP) cells such as double bouquet cells (DBCs) (Chrysanthidis et al., 2019).

### Two-network architecture and connectivity

The network model features two reciprocally connected networks, the so-called Item and Context networks. For simplicity, we assume that Item and Context networks are located at a substantial distance accounting for the reduced between-network connection probabilities (Table 2). Each network follows a cortical architecture with modular structure compatible with previous spiking implementations of attractor memory networks (Lansner, 2009; Tully et al., 2014, 2016; Lundqvist et al., 2011; Fiebig and Lansner, 2017; Chrysanthidis et al., 2019; Fiebig et al., 2020), and is best understood as a subsampled cortical layer 2/3 patch with nested hypercolumns (HCs) and minicolumns (MCs; Fig. 5A). Both networks span a regular spaced grid of 9 HCs (Table 2), each with a diameter of 500 *µ*m (Mountcastle, 1997). In our model, items are embedded in the Item network and context information in the Context network as internal well consolidated long-term memory representations (cell assemblies), supported with within-network weights (within-network connectivity) derived using prior BCPNN long-term learning (Fig. 5B,C). Consequently, these weights were resistant to changes during associative learning of projections between item and context networks (see Results section). Our item and context memory representations are distributed and non-overlapping, i.e. with a single distinct pattern-specific (encoding) MC per HC. This results in a sparse neocortical type of activity patterns (Barth & Poulet 2012). It should be noted that the model tolerates a marginal overlap between different memory patterns, i.e. shared encoding minicolumns (data not shown). Each minicolumn is composed of 30 pyramidal cells (representing the extent of layer 2/3) with shared selectivity, forming a functional (not strictly anatomical) column. In total, the 18 HCs (16 MCs each) of the model contain 8640 excitatory and 1152 inhibitory cells, significantly downsampling the number of MCs per HC (∼100 MCs per HC in biological cortex). Within each HC there are 480 pyramidal cells and 120 basket cells, and hence our model does match in-vivo observations of 4:1 ratio of excitatory to inhibitory cells (Zaitsev and Lewis, 2013). Our model also accounts for another type of inhibition - namely, disynaptic inhibition mediated via dendritic targeting double bouquet and/or bipolar cells. As a result, a sizable fraction of the total inhibition (i.e. all the “learned" inhibition) is modeled implicitly via learned negative weights rather than explicitly via inhibitory cells. The high degree of recurrent connectivity within (Thomson et al., 2002; Yoshimura and Callaway, 2005) and between MCs links coactive MCs into larger cell assemblies (Eyal et al., 2018; Binzegger et al., 2009; Muir et al., 2011; Stettler et al., 2002). Long-range bidirectional between-network connections (item-context bindings or associative connections) are plastic (shown in Fig. 5A only for MC1 in HC1 of the Context network), binding items and contextual information (Ranganath, 2010). On average, recurrent connectivity establishes 100 active plastic synapses onto each pyramidal cell from other pyramidals with the same selectivity, due to a sparse between-network connectivity (*cp_PPA_*) and denser local connectivity (*cp_PP_*, *cp_PPL_*; connection probability refers to the probability that there is a connection between a randomly selected pair of neurons from given populations; in Fig. 6A connection probabilities are only shown for MC1 in HC1 of the Context network). The model yields biologically plausible excitatory postsynaptic potentials (EPSPs) for connections within HCs (0.72 ± 0.085 mV), measured at resting potential *E_L_* (Thomson et al., 2002). Densely recurrent non-specific monosynaptic feedback inhibition mediated by fast spiking inhibitory cells (Kirkcaldie, 2012) implements a local winner-take-all structure (Binzegger et al., 2009) amongst the functional columns. Inhibitory postsynaptic potentials (IPSPs) have an amplitude of -1.160 mV (±0.003) measured at -60 mV (Thomson et al., 2002). These bidirectional connections between basket and pyramidal cells within the local HCs are drawn with a 70% connection probability. Notably, double bouquet cells shown in Figure 6A, are not explicitly simulated, but their effect is nonetheless expressed by the BCPNN rule. A recent study based on a similar single-network architecture (i.e. with the same modular organization, microcircuitry, conductance-based AdEx neuron model, cell count per MC and HC) demonstrated that learned mono-synaptic inhibition between competing attractors is functionally equivalent to the disynaptic inhibition mediated by double bouquet and basket cells (Chrysanthidis et al., 2019). Therefore, BCPNN describes the effect of not-explicitly simulated double-bouquet cells (DBCs) by replacing disynaptic inhibition with negative connections (GABA reversal potential) between cell assemblies that do not share the same pattern selectivity. Also, other network models with negative synaptic weights have been shown to be functionally equivalent to ones with both excitatory and inhibitory neurons with only positive weights (Parisien et al., 2008). Parameters characterizing other neural and synaptic properties including BCPNN can be found in Table 1.

**Figure 6:**
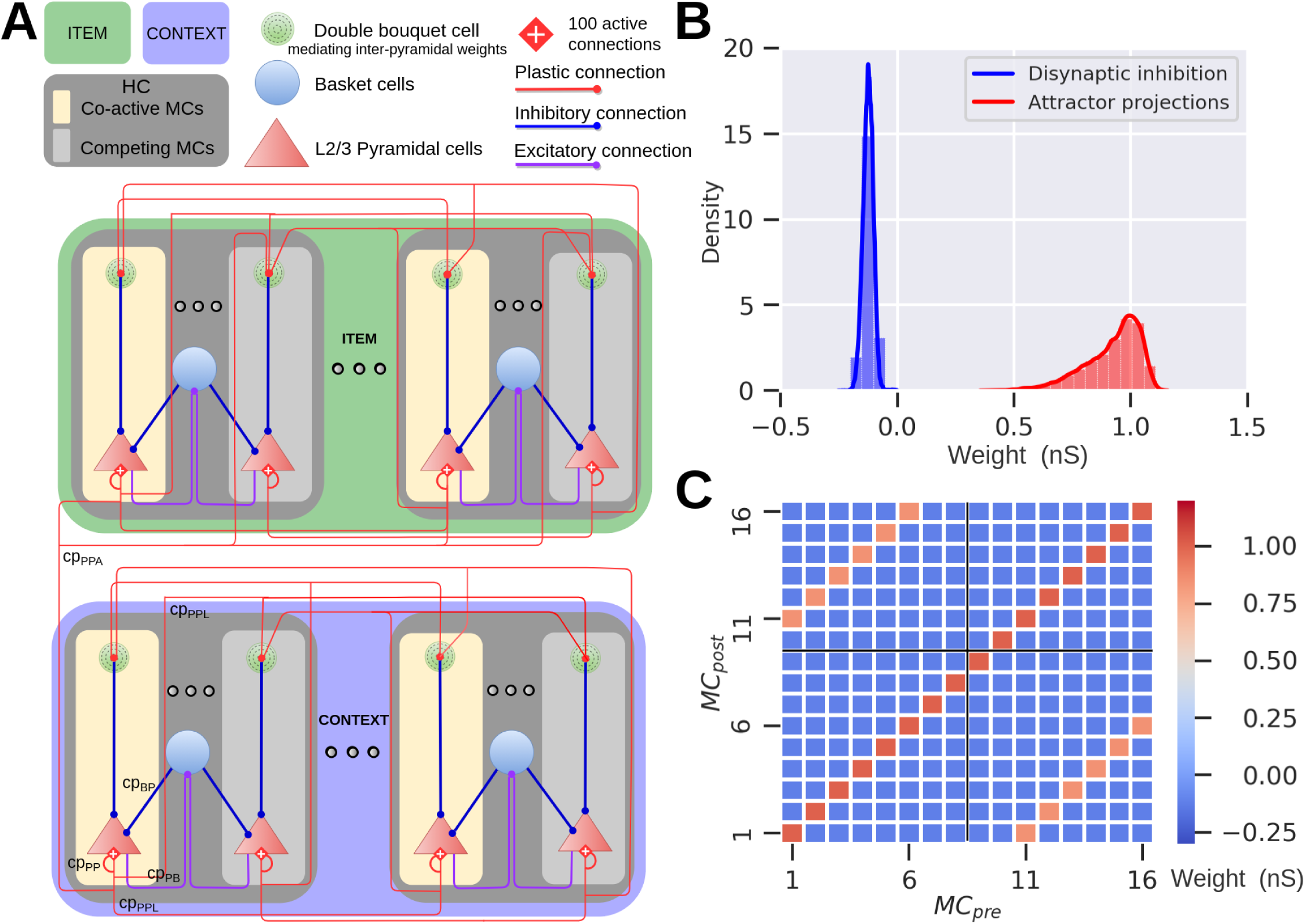
Network architecture and connectivity of the Item (green) and Context (blue) networks. **A**, The model represents a subsampled modular cortical layer 2/3 patch consisting of minicolumns (MCs) nested in hypercolumns (HCs). Both networks contain 9 HCs, each comprising 16 MCs. We preload abstract long-term memories of item and context representations into the respective network, in the form of distributed cell assemblies with weights establishing corresponding attractors. Associative plastic connections bind items with contexts. The network features lateral inhibition via basket cells (purple and blue lines) resulting in a soft winner-take-all dynamics. Competition between attractor memories arises from this local feedback inhibition together with disynaptic inhibition between HCs. **B**, Weight distribution of plastic synapses targeting pyramidal cells. We show the fast AMPA weight components here, but the simulation also includes slower NMDA weight components. **C**, Weight matrix between attractors and competing MCs across two sampled HCs. The matrix displays the mean of the weight distribution between a presynaptic (*MC_pre_*) and postsynaptic minicolumn (*MC_post_*), within the same or different HC (black cross separates grid into blocks of HCs, only two of which are shown here). Recurrent attractor connections within the same HC are stronger (main diagonal, dark red) compared to attractor connections between HCs (off-diagonals, orange). Negative pyramidal-pyramidal weights (blue) between competing MCs amounts to disynaptic inhibition mediated by double bouquet cells.

**Table 2:**
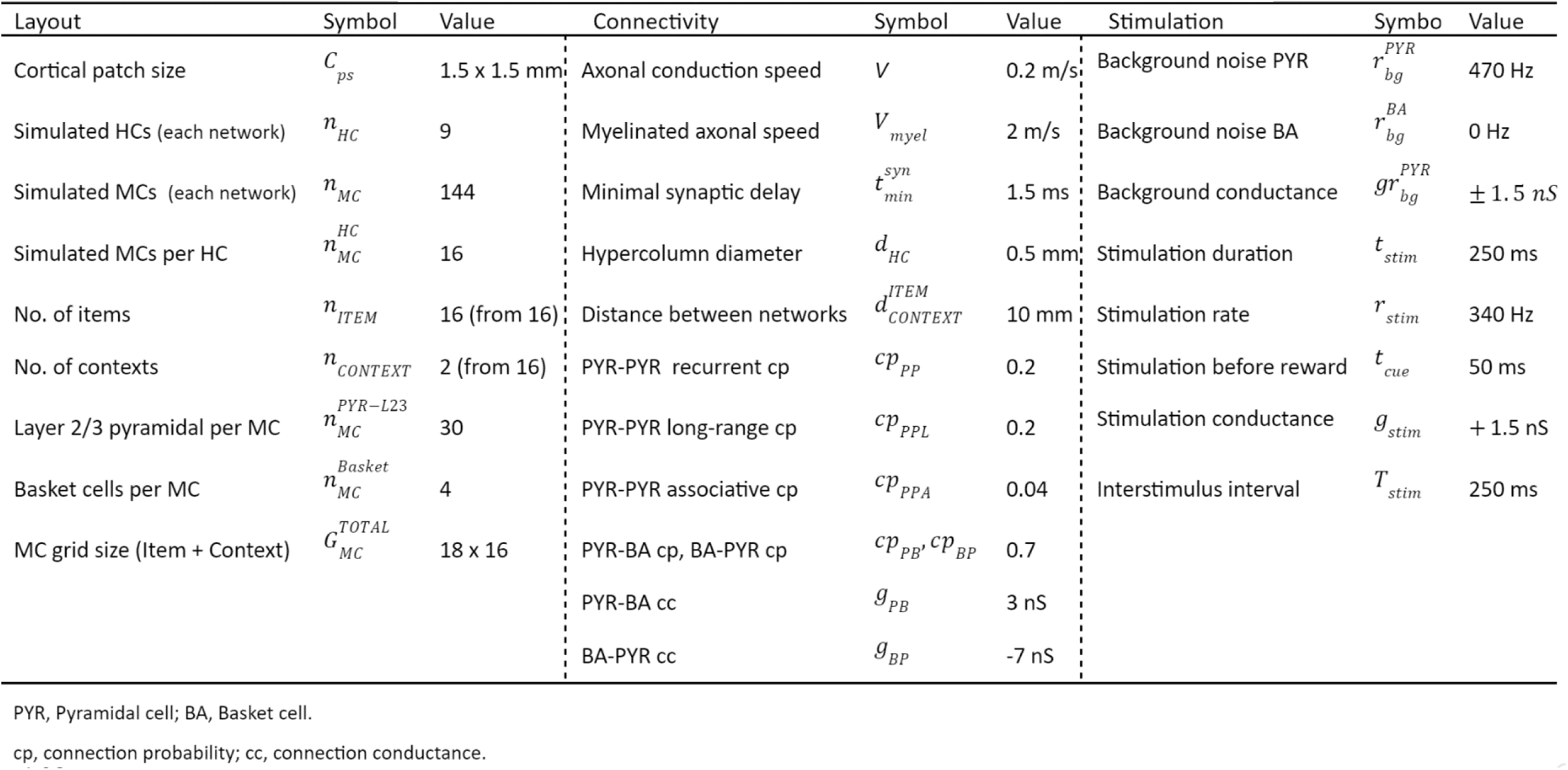
Network layout, connectivity and stimulation protocal.

| Layout | Symbol | Value | Connectivity | Symbol | Value | Stimulation | Symbo | Value |
| --- | --- | --- | --- | --- | --- | --- | --- | --- |
| Cortical patch size | $C_{ps}$ | 1.5 x 1.5 mm | Axonal conduction speed | $V$ | 0.2 m/s | Background noise PYR | $r_{bg}^{PYR}$ | 470 Hz |
| Simulated HCs (each network) | $n_{HC}$ | 9 | Myelinated axonal speed | $V_{myel}$ | 2 m/s | Background noise BA | $r_{bg}^{BA}$ | 0 Hz |
| Simulated MCs (each network) | $n_{MC}$ | 144 | Minimal synaptic delay | $t_{min}^{syn}$ | 1.5 ms | Background conductance | $g_{bg}^{PYR}$ | $\pm 1.5$ nS |
| Simulated MCs per HC | $n_{MC}^{HC}$ | 16 | Hypercolumn diameter | $d_{HC}$ | 0.5 mm | Stimulation duration | $t_{stim}$ | 250 ms |
| No. of items | $n_{ITEM}$ | 16 (from 16) | Distance between networks | $d_{CONTEXT}^{ITEM}$ | 10 mm | Stimulation rate | $r_{stim}$ | 340 Hz |
| No. of contexts | $n_{CONTEXT}$ | 2 (from 16) | PYR-PYR recurrent cp | $cp_{PP}$ | 0.2 | Stimulation before reward | $t_{cue}$ | 50 ms |
| Layer 2/3 pyramidal per MC | $n_{MC}^{PYR-L23}$ | 30 | PYR-PYR long-range cp | $cp_{PPL}$ | 0.2 | Stimulation conductance | $g_{stim}$ | + 1.5 nS |
| Basket cells per MC | $n_{MC}^{Basket}$ | 4 | PYR-PYR associative cp | $cp_{PPA}$ | 0.04 | Interstimulus interval | $T_{stim}$ | 250 ms |
| MC grid size (Item + Context) | $G_{MC}^{TOTAL}$ | 18 x 16 | PYR-BA cp, BA-PYR cp | $cp_{PB}, cp_{BP}$ | 0.7 | | | |
| | | | PYR-BA cc | $g_{PB}$ | 3 nS | | | |
| | | | BA-PYR cc | $g_{BP}$ | -7 nS | | | |

PYR, Pyramidal cell; BA, Basket cell.
cp, connection probability; cc, connection conductance.

Figure 6B shows the weight distributions of embedded distributed cell assemblies, representing different memories stored in the Item and Context networks. Attractor projections can be further categorized into strong local recurrent connectivity within HCs, and slightly weaker long-range excitatory projections across HCs (Fig. 6C).

### Axonal conduction delays

Conduction delays (*t_ij_*) between a presynaptic neuron *i* and a postsynaptic neuron *j* are calculated based on their Euclidean distance, *d*, and a conduction velocity *V* (Eq. 10). Delays are randomly drawn from a normal distribution with a mean according to distance and conduction velocity, with a relative SD of 30% of the mean in order to account for individual arborization differences, and varying conduction speed as a result of axonal thickness and myelination. In addition, a minimal delay of 1.5 ms (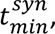, Table 2) is added to reflect synaptic delays due to effects that are not explicitly modeled, e.g. diffusion of neurotransmitters over the synaptic cleft, dendritic branching, thickness of the cortical sheet and the spatial extent of columns (Thomson et al., 2002). Associative between-network projections have a ten-fold faster conduction speed than those within each network, reflecting axonal myelination.

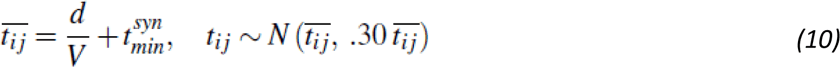

### Stimulation Protocol

Noise input to pyramidal cells is a zero-mean noise, generated by two independent Poisson generators with opposing driving potentials. Pyramidal cells coding for specific items and contexts are stimulated with an additional specific excitation during encoding and cued recall (all parameters in Table 2).

### New- vs old-in-context memory discrimination

Our model discriminates old- vs. new-in-context items based on a comparison of the firing rates in the Item network during stimulation of items in given contexts. New-in-context items were selected if the corresponding trial-average firing rates were 15% lower than the pair-matched old-in-context items. First, we determined a decision threshold high enough to show significant differences between trial-average firing rates, and then we tuned the model (i.e., strength of activations-cues and background excitation - noise) to match the reported behavioral results of an item-in-context memory task. By changing this decision threshold, we can retune the strength of the cues and noise, or even modify other parameters (i.e., boost between-network connectivity) to produce comparable results, so the decision threshold by itself is not critical. While the action selection following the recollection (old item-in-context) was intriguing, it has been out of the scope of this particular study to detail it further.

### Pairwise differences

To show changes in firing rates (f), within- and between-network connectivity (w), and bias (b), we calculate the corresponding differences in the averages between pairs of old and new items, Δf_old-new,_ Δw_old-new,_ Δb_old-new_, respectively.

## Code accessibility

We use the NEST simulator (Gewaltig and Diesmann, 2007), and a custom-built Bayesian-Hebbian learning rule module (BCPNN) in NEST (Tully et al., 2014) running on an HPE Cray EX supercomputer. The spike-based BCPNN learning rule implementation is freely available online at Zenodo (https://doi.org/10.5281/zenodo.5101626). The spiking neural network model is based on an earlier work readily accessible on ModelDB (https://modeldb.science/257610).

## Acknowledgements

This research was supported by Vetenskapsrådet 2018-05360 and 2018-07079, the Swedish e-Science Research Centre (SeRC), Digital Futures, and European Commission, Directorate-General for Communications Networks, Content and Technology (grant no. 101135809). The simulations were performed on resources provided by National Academic Infrastructure for Super­computing in Sweden (NAISS) at the PDC Center for High Performance Computing, KTH Royal Institute of Technology.

## Author contributions

N.C., A.L., and P.H. conceptualized the project; N.C., and P.H. designed research; N.C. performed research; N.C. analyzed data; N.C. curated data; N.C. performed validation; N.C. visualized data; N.C., and F.F. contributed new reagents/analytic tools, N.C. wrote the original draft; N.C., F.F., A.L., and P.H. wrote the paper; P.H acquired funding.

## Supplementary Material

**Figure S1:**
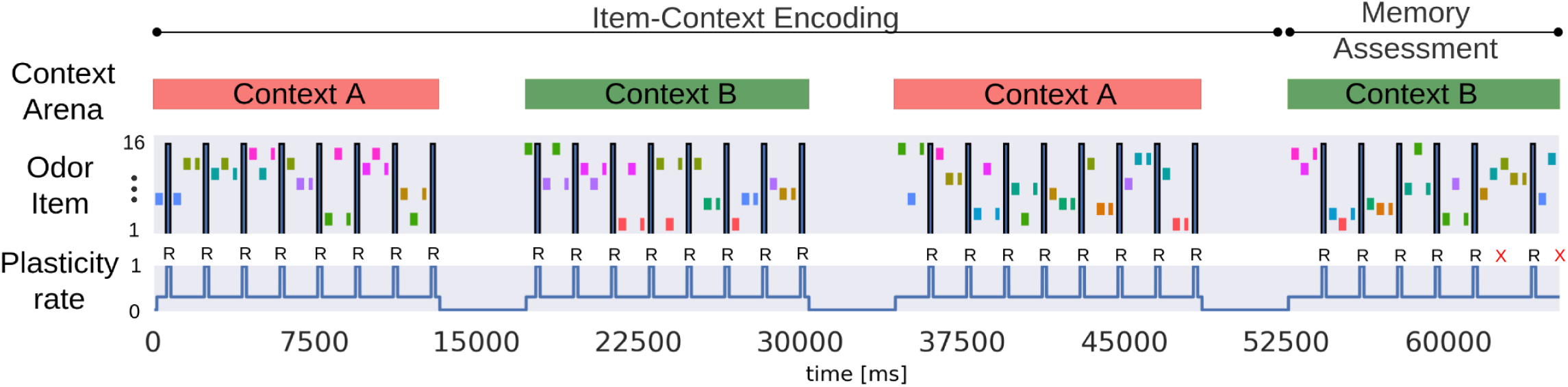
Graphical schematic of the three-context-transition task displaying pairs of new-old odors (depicted as rectangles with unique colors) in a given context. Odors were presented across two contexts in the simulated episodic memory task, and only the new items-in-context were rewarded (**R** symbol in the schematic denotes reward, and **X** symbol, in red, indicates a failed trial) when selected (a 50 ms stimulation of the selected odor preceded the reward phase, representing a final odor sniff before the reward). Once a new item was presented it was considered as old for the subsequent trials in the given context (as a trial we defined a stimulation of a pair of new- and old-in-context items). Items were stimulated for the first time in context A, half of the total 16 items were presented and rewarded in context A. After the context transition half of the 16 items were presented in random pairs in context B. After one more context transition, we activated in context A the remaining 8 items that were not previously presented in that context. Finally, Memory Assessment was made in context B, where we presented the remaining half of the items that had not been presented in context B, and paired them randomly with old items (pairs of odors were different throughout the task). Context representations were constantly activated while cueing pairs of new-old items for 250 ms each. In the Memory Assessment block, pairs of new-old items followed the Arrangement 1 criterion (new items were encoded more recently than the old ones). While context representations were persistently cued we activated new and old items-in-context during trials. Plasticity rate of the associative binding between Item and Context networks was modulated during item presentation and rewarded accordingly (bottom subplot).

**Figure S2:**
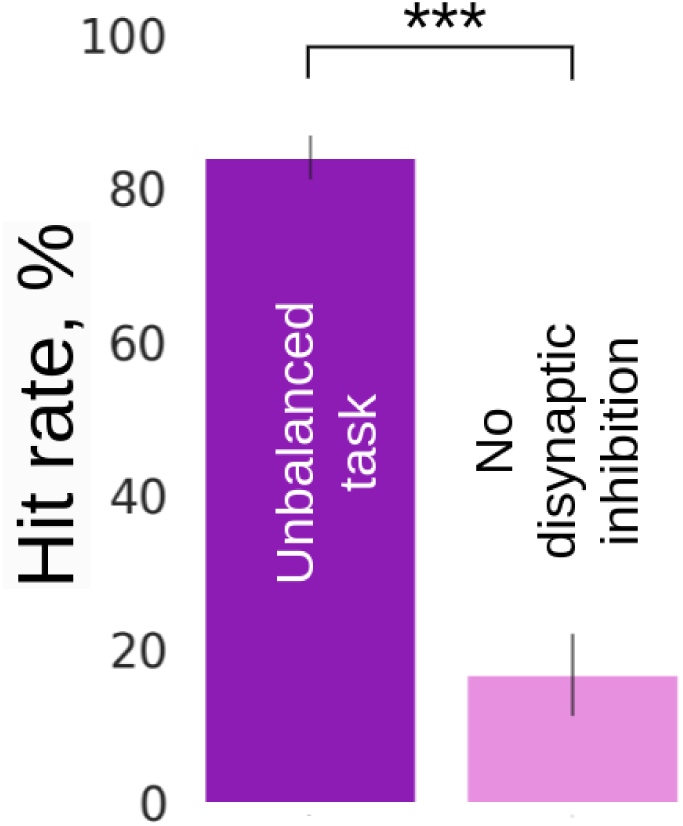
Average recall performance (hit rate, %) for the unbalanced and “No disynaptic inhibition” prediction tasks corresponding to Arrangement 1 configuration. For the No disynaptic inhibition task, the inhibitory weights between networks were disabled, and thus the Context network did not suppress new items during Memory Assessment. SDs derived from the Bernoulli distributions for the probabilities of success (hit) across all trials (scaled to %).

## Notes

### Competing Interest Statement

The authors have declared no competing interest.

## References

Barth AL, Poulet JF (2012) Experimental evidence for sparse firing in the neocortex. Trends Neurosci 35:345–355.

Bevins, R. A., and Besheer, J. (2006). Object recognition in rats and mice: A one-trial non-matching-to-sample learning task to study ‘recognition memory’. Nat. Protoc. 1 (3), 1306–1311. doi:10.1038/nprot.2006.205

Binzegger T, Douglas RJ, Martin KA (2009) Topology and dynamics of the canonical circuit of cat v1. Neural Netw 22:1071–1078.

Brea J, Clayton NS, Gerstner W. Computational models of episodic-like memory in food-caching birds. Nature Communications. 2023; 14(1):2979. 10.1038/s41467-023-38570-x PMID: 37221167

Brette R, Gerstner W (2005) Adaptive exponential integrate-and-fire model as an effective description of neuronal activity. J Neurophysiol 94:3637–3642.

Caporale N, Dan Y (2008) Spike timing–dependent plasticity: a Hebbian learning rule. Annu Rev Neurosci 31:25–46.

Chrysanthidis, N., Fiebig, F., Lansner, A., 2019. Introducing double bouquet cells into a modular cortical associative memory model. J Comput Neurosci 47, 223–230. 10.1007/s10827-019-00729-1

Chrysanthidis, N., Fiebig, F., Lansner, A., Herman, P., 2022. Traces of Semantization, from Episodic to Semantic Memory in a Spiking Cortical Network Model. eNeuro 9, ENEURO.0062-22.2022. 10.1523/ENEURO.0062-22.2022

DeFelipe, J., Ballesteros-Yáñez, I., Inda, M.C., Muñoz, A. (2006). Double-bouquet cells in the monkey and human cerebral cortex with special reference to areas 17 and 18. Progress in Brain Research, 154, 15–32.

Ecker, U. K. H., Zimmer, H. D., Groh-Bordin, C., & Mecklinger, A. (2007). Context effects on familiarity are familiarity effects of context—An electrophysiological study. International Journal of Psychophysiology: Official Journal of the International Organization of Psychophysiology, 64(2), 146–156. 10.1016/j.ijpsycho.2007.01.005

Erickson MA, Maramara LA, Lisman J: A single brief burst induces GluR1-dependent associative short-term potentiation: a potential mechanism for short-term memory. J Cognit Neurosci 2010:2530–2540, 10.1162/jocn.2009.21375

Eyal G, Verhoog MB, Testa-Silva G, Deitcher Y, Benavides-Piccione R, DeFelipe J, De Kock CP, Mansvelder HD, Segev I (2018) Human cortical pyramidal neurons: from spines to spikes via models. Front Cell Neurosci 12:181.

Fiebig F, Lansner A (2017) A spiking working memory model based on Hebbian short-term potentiation. J Neurosci 37:83–96.

Fiebig F, Herman P, Lansner A (2020) An indexing theory for working memory based on fast Hebbian plasticity. eNeuro 7:ENEURO.037419.2020.

Gerstner W, Naud R (2009) How good are neuron models? Science 326:379–380.

Gewaltig MO, Diesmann M (2007) Nest (neural simulation tool). Scholarpedia 2:1430.

Greve, A., Donaldson, D.I., van Rossum, M.C.W., 2009. A single-trace dual-process model of episodic memory: A novel computational account of familiarity and recollection. Hippocampus NA-NA. 10.1002/hipo.20606

Groh-Bordin, C., Zimmer, H.D., Ecker, U.K.H., 2006. Has the butcher on the bus dyed his hair? When color changes modulate ERP correlates of familiarity and recollection. NeuroImage 32, 1879– 1890. 10.1016/j.neuroimage.2006.04.215

Hayes, S. M., Nadel, L., & Ryan, L. (2007). The effect of scene context on episodic object recognition: Parahippocampal cortex mediates memory encoding and retrieval success. Hippocampus, 17(9), 873–889. 10.1002/hipo.20319

Herman, P.A., Lundqvist, M., Lansner, A., 2013. Nested theta to gamma oscillations and precise spatiotemporal firing during memory retrieval in a simulated attractor network. Brain Research, Selected papers presented at the Tenth International Neural Coding Workshop, Prague, Czech Republic, 2012 1536, 68–87. 10.1016/j.brainres.2013.08.002

Kanatsou S., Krugers H., Object-context recognition memory test for mice, Bioprotocol 6 (2016) e1925, 10.21769/Bio.Protoc.1925.

Kelsom, C., & Lu, W. (2013). Development and specification of GABAergic cortical interneurons. Cell & Bioscience, 3(1), 19.

Krimer, L.S., Zaitsev, A.V., Czanner, G., Kroner, S., González-Burgos, G., Povysheva, N.V., Iyengar, S., Barrionuevo, G., Lewis, D.A. (2005). Cluster analysis–based physiological classification and morphological properties of inhibitory neurons in layers 2–3 of monkey dorsolateral prefrontal cortex. Journal of Neurophysiology, 94(5), 3009–3022.

Lansner A, Ekeberg Ö (1989) A one-layer feedback artificial neural network with a Bayesian learning rule. Int J Neur Syst 01:77–87.

Lansner A (2009) Associative memory models: from the cell-assembly theory to biophysically detailed cortex simulations. Trends Neurosci 32:178–186.

Lesburguères, E., Tsokas, P., Sacktor, T., & Fenton, A. (2017). The Object Context-place-location Paradigm for Testing Spatial Memory in Mice. BIO-PROTOCOL, 7(8). 10.21769/BioProtoc.2231

Lisman J (2017). Glutamatergic synapses are structurally and biochemically complex because of multiple plasticity processes: long-term potentiation, long-term depression, short-term potentiation and scaling. Philos Trans R Soc Lond B Biol Sci 2017, 10.1098/rstb.2016.0260

Lu, Q., Hasson, U. & Norman, K. A. A neural network model of when to retrieve and encode episodic memories. eLife 11, e74445 (2022).

Lundqvist M, Herman P, Lansner A (2011) Theta and gamma power increases and alpha/beta power decreases with memory load in an attractor network model. J Cogn Neurosci 23:3008–3020. 10.1162/jocn_a_00029

Martin-Ordas, G., Atance, C.M., Caza, J.S., 2014. How do episodic and semantic memory contribute to episodic foresight in young children? Front Psychol 5, 732. 10.3389/fpsyg.2014.00732

Merkow, M.B., Burke, J.F., Kahana, M.J., 2015. The human hippocampus contributes to both the recollection and familiarity components of recognition memory. Proceedings of the National Academy of Sciences 112, 14378–14383. 10.1073/pnas.1513145112

Mongillo, G., Barak, O., Tsodyks, M., 2008. Synaptic Theory of Working Memory. Science 319, 1543– 1546. 10.1126/science.1150769

Mountcastle VB (1997) The columnar organization of the neocortex. Brain: a journal of neurology 120:701–722.

Muir DR, Da Costa NM, Girardin CC, Naaman S, Omer DB, Ruesch E, Grinvald A, Douglas RJ (2011) Embedding of cortical representations by the superficial patch system. Cereb Cortex 21:22442260.

Norman, K.A., O’Reilly, R.C., 2003. Modeling hippocampal and neocortical contributions to recognition memory: A complementary-learning-systems approach. Psychological Review 110, 611–646. 10.1037/0033-295X.110.4.611

O’Brien, J., & Sutherland, R. J. (2007). Evidence for episodic memory in a pavlovian conditioning procedure in rats. Hippocampus, 17(12), 1149–1152. 10.1002/hipo.20346

Panoz-Brown, D., Corbin, H.E., Dalecki, S.J., Gentry, M., Brotheridge, S., Sluka, C.M., Wu, J.-E., Crystal, J.D., 2016. Rats Remember Items in Context Using Episodic Memory. Current Biology 26, 2821–2826. 10.1016/j.cub.2016.08.023

Parisien, Christopher and Anderson, Charles H and Eliasmith, Chris (2008) Solving the problem of negative synaptic weights in cortical models. Neural computation 6: 1473–1494

Peters, A., & Yilmaz, E. (1993). Neuronal organization in area 17 of cat visual cortex. Cerebral Cortex, 3, 49-68.

Ren Q, Kolwankar KM, Samal A, Jost J (2010) Stdp-driven networks and the c. elegans neuronal network. Physica A: Statistical Mechanics and its Applications 389:3900–3914.

Sederberg, P. B., Howard, M. C., & Kahana, M. J. (2008). A context-based theory of recency and contiguity in free recall. Psychological Review, 115, 893–912.

Stettler DD, Das A, Bennett J, Gilbert CD (2002) Lateral connectivity and contextual interactions in macaque primary visual cortex. Neuron 36:739–750.

Tam, S. K. E., Bonardi, C. & Robinson, J. (2015) Relative recency influences object-in-context memory. Behav. Brain Res. 281, 250–257.

Thomson AM, West DC, Wang Y, Bannister AP (2002) Synaptic connections and small circuits involving excitatory and inhibitory neurons in layers 2–5 of adult rat and cat neocortex: triple intracellular recordings and biocytin labelling in vitro. Cereb Cortex 12:936953.

Tsodyks MV, Markram H (1997) The neural code between neocortical pyramidal neurons depends on neurotransmitter release probability. Proc Natl Acad Sci U S A 94:719–723.

Tully, P.J., Hennig, M.H., Lansner, A., 2014. Synaptic and nonsynaptic plasticity approximating probabilistic inference. Front. Synaptic Neurosci. 6. 10.3389/fnsyn.2014.00008

Tully PJ, Lindén H, Hennig MH, Lansner A (2016) Spike-based Bayesian-Hebbian learning of temporal sequences. PLoS Comput Biol 12:e1004954.

Tulving E (1972) 12. episodic and semantic memory. Organization of memory/Eds E. Tulving, W. Donald- son, NY: Academic Press pp. 381–403.

Tulving, E., 1985. Memory and consciousness. Canadian Psychology/Psychologie canadienne 26, 1–12. 10.1037/h0080017

Van Rossum MC, Bi GQ, Turrigiano GG (2000) Stable hebbian learning from spike timing-dependent plasticity. Journal of neuroscience 20:8812–8821.

Umanath, S., Coane, J.H., 2020. Face Validity of Remembering and Knowing: Empirical Consensus and Disagreement Between Participants and Researchers. Perspect Psychol Sci 15, 1400–1422. 10.1177/1745691620917672

Wahlgren N, Lansner A (2001) Biological evaluation of a HebbianBayesian learning rule. Neurocomputing 38–40:433–438.

Wang, Y., Markram, H., Goodman, P.H., Berger, T.K., Ma, J., Goldman-Rakic, P.S., 2006. Heterogeneity in the pyramidal network of the medial prefrontal cortex. Nature neuroscience 9, 534–42. 10.1038/nn1670

Ward, G., & Tan, L. (2023). The role of rehearsal and reminding in the recall of categorized word lists. Cognitive Psychology, 143, 101563.

Weisz, V. I., Rios, M. B., & Argibay, P. F. (2012). Episodic-like memory: New perspectives from a behavioral test in rats. Journal of Integrative Neuroscience, 11(01), 1–15. 10.1142/S021963521250001X

Wilson DI, Langston RF, Schlesiger MI, Wagner M, Watanabe S, Ainge JA. 2013 Lateral entorhinal cortex is critical for novel object-context recognition. Hippocampus 23, 352– 366. (doi:10.1002/hipo.22095)

Wixted, J.T., 2007. Dual-process theory and signal-detection theory of recognition memory. Psychological Review 114, 152–176. 10.1037/0033-295X.114.1.152

Yonelinas AP, Aly M, Wang WC, Koen JD (2010) Recollection and familiarity: Examining controversial assumptions and new directions. Hippocampus 20:1178–1194. 10.1002/hipo.20864

Yonelinas, A.P., Ranganath, C., Ekstrom, A.D., Wiltgen, B.J., 2019. A contextual binding theory of episodic memory: systems consolidation reconsidered. Nat Rev Neurosci 20, 364–375. 10.1038/s41583-019-0150-4

Yoshimura Y, Callaway EM (2005) Fine-scale specificity of cortical networks depends on inhibitory cell type and connectivity. Nat Neurosci 8:1552–1559.

Zaitsev, A.V., Lewis, D.A., 2013. Functional properties and short-term dynamics of unidirectional and reciprocal synaptic connections between layer 2/3 pyramidal cells and fast-spiking interneurons in juvenile rat prefrontal cortex. Eur. J.Neurosci. 38, 2988e2998.

Zhang, K., Bromberg-Martin, E. S., Sogukpinar, F., Kocher, K., & Monosov, I. E. (2022). Surprise and recency in novelty detection in the primate brain. Current Biology, 32(10), 2160–2173.e6. 10.1016/j.cub.2022.03.064

